# Mapping kinase-dependent tumor immune adaptation with multiplexed single-cell CRISPR screens

**DOI:** 10.64898/2026.01.08.698516

**Authors:** Lingting Shi, Ross M. Giglio, Qingyuan Cai, Mathini Vaikunthan, Justin Hong, Abdullah Naqvi, Marina Milea, Hannah Khanshali, Anna Schoonen, Nicholas Hou, Jonathan Guo, Melanie Fraidenburg, Xumin Shen, Seth W. Malinowski, Keith L. Ligon, Raúl Rabadán, Elham Azizi, José L. McFaline-Figueroa

## Abstract

Immune dysfunction in cancer is enacted by multiple programs, including tumor cell-intrinsic responses to distinct immune subpopulations. A subset of these immune evasion programs can be systematically recapitulated through direct tumor-immune interactions *in vitro*. Here, we present an integrated, high-throughput single-cell CRISPR screening framework focused on the protein kinome for mapping the tumor-intrinsic regulation of T cell-driven immune pressure in glioblastoma (GBM). We combine pooled CRISPR interference and activation (CRISPRi/a) with immune-matched NY-ESO-1 antigen-specific allogeneic GBM-T cell co-culture and massively multiplexed single-cell transcriptomics to systematically quantify how genetic perturbation reshapes baseline tumor state and adaptive responses across graded effector-to-target ratios. We further leverage deep generative models for analyzing pooled CRISPR screens to decipher the effects of genetic perturbations on the mechanisms of tumor resistance. This framework resolves distinct modules of immune evasion and survival, including the regulation of the antigen-presentation machinery, interferon/NF-κB signaling, oxidative stress resilience, and checkpoint/cytokine programs, while identifying perturbations that reroute the continuous tumor transcriptional trajectory induced by T cell engagement. A secondary chemical screen in patient-derived GBM cultures identified putative kinase targets of immune evasion phenotypes (e.g., EPHA2 and PDGFRA), whose inhibition leads to the blockade of evasive programs and enhances T cell-mediated GBM killing. Together, this workflow provides a scalable blueprint for comprehensive charting of the genetic control of tumor-immune interactions.

## Introduction

The clinical success of immunotherapy has fundamentally shifted the oncology landscape, yet the majority of patients with solid tumors fail to achieve durable responses^1^. Effective anti-tumor immunity requires sustained activation and effector function of cytotoxic T cells. However, tumors have evolved multiple mechanisms to evade immune destruction, operating both at a distance through soluble factors (TGF-β^2^, IL-10^3^) and locally through direct physical interactions with infiltrating immune cells. The most clinically relevant examples are checkpoint pathways like PD-1/PD-L1 and CTLA-4^4^ that engage in direct tumor-T cell contact to deliver inhibitory signals. The composition and deployment of immune evasion mechanisms varies substantially across cancer types, shaped by tissue context, mutational landscape, and the tumor microenvironment^5–7^. Despite extensive characterization of individual pathways, how tumor-intrinsic programs are dynamically engaged by direct tumor-immune interactions remains incompletely understood.

Upon interacting with cancer cells, activated T cells release IFN-γ^8^ that induces upregulation of PD-L1 on tumor cells^9^. PD-L1 then engages PD-1 on T cells, suppressing their activation, proliferation, and effector function^10^, while maintaining Treg phenotypes^11,12^. In some cancer types, upregulation of PD-L1 has been associated with more aggressive disease and poorer clinical outcomes (e.g., hepatocellular carcinoma^13^, breast cancer^14^, gastric cancer^15^, prostate cancer^16^, glioblastoma multiforme (GBM)^17^). Thus, checkpoint inhibitors (PD-1/PD-L1) and cytokine therapy have been explored to reverse cytotoxic T cell dysfunction. The antigen processing and presentation pathway is also crucial for an efficient response to immune checkpoint inhibitor therapy^18^. Numerous cancer types frequently downregulate components of antigen processing and presentation machinery (APM)^19–21^ as a mechanism of immune evasion. Overcoming these barriers, adaptive immune checkpoint signaling coupled with defective antigen presentation, represents a critical unmet challenge for restoring effective anti-tumor immunity.

Glioblastoma multiforme (GBM) represents an extreme manifestation of these immune evasion strategies, providing a compelling model to study tumor-intrinsic regulation of anti-tumor immunity. Despite an intense treatment regimen of radiation, chemotherapy, and tumor-treating fields, most patients succumb to the disease within 14-15 months of diagnosis, highlighting the dire need for more effective therapies^22^. GBM is considered an immunologically "cold" tumor that is particularly resistant to immunotherapy^23–26^. Nevertheless, T cell infiltration is detectable in primary tumors^27^, where infiltrating T cells exhibit a more GZMK phenotype that does not express high levels of classic cytotoxic markers such as *GZMB*, *IFNG*, and *PRF1^28,29^*. Notably, robust T cell infiltration is also observed both in recurrent GBM^30,31^ and in the first clinical trial of EGFR-targeting CAR T cells in glioblastoma^32^. Moreover, emerging clinical evidence demonstrates that the subset of GBM patients who do respond to immunotherapy can achieve durable, lasting responses^33,34^, suggesting that tumor-intrinsic mechanisms may be therapeutically reprogrammed. These observations underscore the critical need to identify the tumor-intrinsic factors that modulate GBM immunogenicity and could be targeted to render tumors more susceptible to immune attack. At the same time, the availability of well-defined antigen-specific T cell models and patient-derived cultures makes GBM a tractable system for scalable, mechanistic perturbation studies.

Recently, single-cell CRISPR-based genetic screens were used to define the regulation driving immunogenicity in melanoma at the level of T cell-mediated tumor killing^35^, providing targets that could decrease immune evasion. In glioblastoma, Larson et al.^36^ performed a genome-wide CRISPR screen in U87 cells co-cultured with EGFR-targeted CAR T cells and identified the IFN-γ receptor pathway as a critical regulator of CAR T-mediated killing in solid tumors. However, existing studies have largely focused on binary survival readouts or limited immune contexts and have not systematically resolved how tumor-intrinsic programs are transcriptionally reconfigured across graded immune pressure at single-cell resolution. Such approaches are constrained by challenges in creating scalable model systems for screening and our incomplete understanding of brain immunity, making it unclear which factors to prioritize.

Kinases play a crucial role in various cellular processes regulating a wide range of cellular activities^37^, including signal transduction and the cell cycle^38^. Dysregulation of kinase signaling is common in cancer. Fortunately, kinases are largely druggable targets, with 80 FDA-approved therapeutic agents currently available^39^. Giglio et al., previously identified EGFR inhibitors that can potentially modulate APM expression as well as T cell-mediated killing^40^. In addition, Wang et al., have shown that combining immunotherapy and kinase inhibitors (e.g., ATR) improves GBM survival rates due to increased CD8^+^ T cell infiltration^41^. More broadly, protein kinase inhibitors, demethylating drugs, and immunomodulatory drugs have all been reported to exhibit potential synergy in combination with adoptive cell therapy (CAR-T)^42^. Understanding how tumor-intrinsic kinase networks orchestrate immune evasion during direct T cell engagement can be leveraged with immunotherapy to make GBM more immunogenic. Thus, in this study, we systematically interrogated GBM-T cell interactions across the entire human kinome using genetic perturbations coupled with high-throughput single-cell sequencing to identify tumor-intrinsic immunomodulators that may inform rational combinatorial treatment designs.

In this study, we established a scalable, immune-matched GBM-T cell co-culture platform using the NY-ESO-1 antigen system to systematically interrogate tumor-immune interactions. By integrating pooled genetic perturbations (CRISPRi and CRISPRa) with highly multiplexed single-cell profiling using sci-Plex-GxE^43^, we generated a quantitative, single-cell map of kinase-dependent immune modulation across graded cytotoxic stress. Leveraging deep learning-based analytical approaches, we identified a set of candidate immunomodulatory kinases, which we subsequently validated using small-molecule inhibitors in patient-derived GBM neurospheres within the same NY-ESO-1 antigen framework. Notably, inhibition of PDGFRA and EPHA2 enhanced T cell-mediated cytotoxicity, revealing actionable tumor-intrinsic resistance mechanisms. Together, these results nominate PDGFRA and EPHA2 as promising combinatorial targets to potentiate immunotherapy in glioblastoma.

## Results

### Increasing T cell exposure reshapes tumor cell state toward immune resistance

To map the crosstalk between glioblastoma (GBM) and cytotoxic T cells, we co-cultured NY-ESO-1-expressing GBM cells with allogeneic T cells at increasing effector-to-target (E:T) ratios (Figure 1a, Supplementary Figure 1). An effector-to-target ratio of 2:1 (T cell:GBM) induced robust tumor cell killing within 18 hours (Supplementary Figure 1d). Gene expression profiling of tumor-immune co-cultures revealed a ratio-dependent transcriptional response (Figure 1b, Supplementary Figure 2), hereafter referred to as the T cell-induced program (TCIP), which captures tumor-intrinsic transcriptional adaptation to increasing immune pressure. Analysis of the top 30 differentially expressed genes highlighted a coordinated induction of inflammatory signaling, stress adaptation, and tumor-immune interaction programs (Figure 1c,d). These included activation of NF-κB and cytokine signaling (*NFKB1*, *TNFAIP3*, *CSF3*, *CXCL2*, *EREG*)^44–47^, interferon-associated immune response genes (*STAT1*, *GBP1*, *ICAM1*, *CD74*)^48–50^, and metabolic and oxidative stress regulators (*SOD2*, *NAMPT*, *KYNU*, *MT2A*)^51–54^, consistent with tumor-intrinsic responses to increasing T cell pressure.

**Figure 1.**
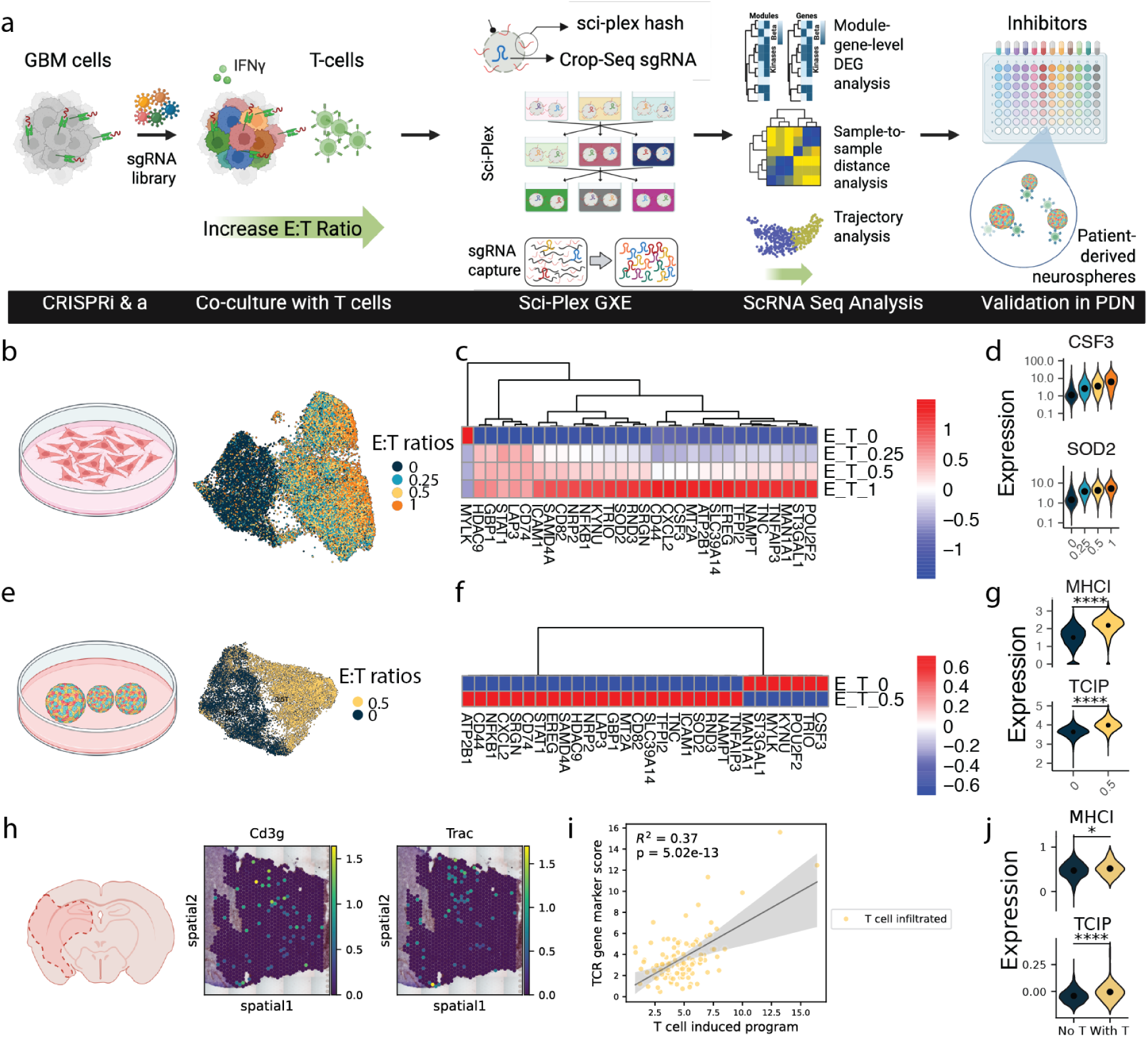
Cytotoxic T cells induce dynamic changes in gene expression including increases in factors associated with immune evasion. (a) Experimental workflow. NY-ESO-1-expressing glioblastoma (GBM, U87) cells were transduced with a pooled CRISPRi/a kinome library (∼500 kinases) and co-cultured with immune-matched, antigen-specific CD8+ T cells across graded effector-to-target (E:T) ratios (effector: T cell, target: GBM). Following an 18-hour exposure, tumor cells were processed with sci-Plex for multiplexed single-cell RNA-seq and sgRNA capture. Transcriptomes were analyzed using generalized linear modeling (GLM) for gene-level perturbation effects, MrVI for sample-level heterogeneity, and Decipher for trajectory analysis across E:T ratios, followed by small-molecule validation in patient-derived neurospheres. (b) U87 and T cell co-culture: UMAP visualizations, generated from PCA space. (c) A representative heatmap highlights perturbation-specific shifts in modules such as oxidative stress (*SOD2*) and cytokine signaling (*CSF3*). (d) Violin plots showing *CSF3* and *SOD2* expression across increasing effector:target (E:T) ratios. (e) The UMAP, generated from PCA space, shows distinct separation of GBM cells with versus without T cell exposure from patient derived neurospheres. (f) A representative heatmap highlights perturbation-specific shifts in modules such as oxidative stress (*SOD2*) and cytokine signaling (*CSF3*). (g) Violin plots showing MHC-I pathways and T cell induced program (TCIP) scores in co-cultured GBM cells. (h) *Cd3g*, and *Trac* gene expression of murine glioma tissue. (i) Scatter plot illustrates correlations between T cell infiltration represented by TCR gene marker score level and T cell-induced program genes. (j) Violin plots showing MHC-I pathways and T cell induced program genes on tissue spots with or without T cell infiltration. TCIP, T cell-induced program. Statistical tests were performed two-sided using the Mann-Whitney U test (*p<0.05, and ****p<0.0001).

Extending this analysis to a broader set of differentially expressed genes clarified the directionality of the underlying cell-state transition (Supplementary Table 1). We found upregulation of pro-inflammatory cytokines and chemokines (*CXCL8*, *CXCL2, CXCL3*, *CXCL9*, *CXCL11*, *CSF3* (G-CSF), *IL1B*, and *IL32*)^55,56^, IFN-γ-induced and antigen-presentation-associated genes (GBPs, *IRF1*, and *BATF2*)^57,58^, NF-κB pathways (*NFKBIA*, *NFKBIZ*, *NFKB1*)^59^, immune checkpoint (*CD274* (PD-L1), *PDCD1LG2* (PD-L2), *IDO1*)^60–62^, and stress response, apoptosis resistance & survival (*SOD2*, *NAMPT*, *BIRC3*, *IER3*)^53,54,63,64^. In contrast, genes associated with progenitor-like and developmental states (*DLX2*, *DLX3*, *SOX4*, *SOX9*)^65,66^, stemness (*ID1*, *ID3*)^67^, and migratory, adhesion, and mechanical sensing programs (*EPHA3*, *MYLK*, *EPB41L1*, *RAP2B*)^68–71^ were downregulated under immune pressure. Together, these changes indicate a cell-state transition from a plastic, migratory, stem-like tumor state toward an inflammatory, stress-adapted, and immune-resistant state with increasing T cell exposure. Both IFN-γ and NF-κB signaling pathways have been identified as core regulators for tumor-intrinsic cytotoxic T lymphocyte evasion^72^. NF-κB signaling, in particular, has been explored as a therapeutic target in cancer immunotherapy and has been shown to directly regulate the expression of immune checkpoint molecules^73^.

To determine whether these phenotypes obtained *in vitro* are conserved in patient-derived models, we similarly overexpressed NY-ESO-1 antigen as well as HLA-A*02:01 to immune match to NY-ESO-1 TCR T cells in patient-derived neurosphere (PDN) lines (Figure 1a,e). PDNs are composed of glioma stem-like cells (GSCs) that self-renew, differentiate into multiple lineages, and recapitulate the heterogeneity of the original tumor, providing a more clinically relevant platform for studying tumor formation and therapeutic resistance. For PDN models, we observed induction of a similar set of T cell-induced program signatures and MHC class I-related complexes following T cell exposure (Figure 1f,g).

To further determine whether these changes are also observed as a function of tumor-immune interactions *in vivo* we analyzed spatial transcriptomics data from the GL261 mouse model from García-Vicente et al.,^74^ (Figure 1h). Spatial analysis revealed that TCR transcripts (*Trac*, *Trbc1*, and *Trbc2*), marking infiltrating T cells, correlated with elevated expression of T cell-induced program genes (Figure 1i). Tumor regions containing T cells further showed increased expression of these genes and enhanced MHC class I compared to regions lacking T cell infiltration (Figure 1j).

Co-culturing NY-ESO-1-expressing GBM cells with autologous T cells revealed a ratio-dependent transcriptional response in GBM cells characterized by the induction of inflammatory, stress-adaptation, and cell-interaction programs, which was conserved in patient-derived neurosphere models and observed *in vivo* in a mouse model.

### Cancer-T Cell Interactions Drive T Cell Effector Remodeling and Bidirectional Cytotoxic-Immunosuppressive Signaling

Cancer cells also induce pronounced transcriptomic changes in T cells (Supplementary Figure 3a,b). Analysis of the T cell compartment of our co-cultures revealed four distinct T cell states (clusters). Cluster 0 was enriched for T cells that had not been exposed to cancer cells and was characterized by high expression of *GZMA*, *GNLY*, and *IL7R* (Supplementary Figure 3c,d). In contrast, Cluster 1, which was dominated by T cells exposed to cancer cells, exhibited upregulation of cytotoxic and activation markers, including *GZMB* and *IL2RA*. The top TCR clonotypes were also identified by enriching for TCR CDR3 sequences from 3’ 10X cDNA library with PCR (see Methods, Supplementary Figure 3e).

We used nichenet^75^ to examine T cell-GBM crosstalk via ligand-receptor interaction analysis identifying crosstalk associated with cytotoxic signaling pathways and immunosuppressive mechanisms that shape the tumor microenvironment (Supplementary Figure 3f). Tumor necrosis factor (TNF) family ligands were upregulated by T cells engaging their corresponding receptors on GBM cells (*TNFRSF1A, TNFRSF1B*, *TNFRSF21*)^76^. TNF signaling promotes programmed cell death (apoptosis) in GBM, representing a cytotoxic mechanism employed by T cells upon cancer cell engagement^76^. In contrast, GBM cells express multiple immunosuppressive ligands that counteract T cell anti-tumor responses. *PVR* (poliovirus receptor) expressed by GBM engages inhibitory *CD96* receptors on T cells that reduce their cytotoxic response^77,78^. *LAMA5* from GBM cells interacts with multiple integrin receptors (*ITGA2*, *ITGA6*, *ITGB1*) on T cells, which could be relevant to adhesion and activation. These results further highlight the complexity of tumor-immune interactions that occur within the context of our allogeneic models.

### Kinase perturbation alters antigen presentation and immune evasion programs

Building on these observations, we next aimed to systematically characterize how kinase perturbations reshape glioblastoma responses to T cell exposure (Figure 2a). We genetically perturbed NY-ESO-1 overexpressing GBM cells to express a system for CRISPR-mediated gene inactivation (CRISPRi: dCas9-KRAB) or overexpression (CRISPRa: dCas9-SunTag) by delivering CRISPR single-guide RNAs targeting all human protein kinases and co-cultured tumor and immune cells at increasing effector:target ratios (Supplementary Figure 4,5). Embedding of cells in a learned transcriptional manifold again revealed progressive transcriptional divergence at increasing E:T ratios, reflecting the ratio-dependent impact of T cell exposure (Figure 2b, Supplementary Table 3). Examining the consequence of kinase perturbation on known immune-associated pathways revealed kinase specific modulation of the antigen-processing and presentation (APM) and inflammatory pathways including kinase modulators of MHC class I, MHC class II, interferon-γ (IFN-γ), and NF-κB programs many of which, showed differential transcriptional shifts as a function of gene knockdown and overexpression (Figure 2c). While some kinases broadly influenced the APM (e.g., *ERBB2,IRAK3*, *TLK1, EGFR (Supplementary Table 3)*), others selectively affected interferon or NF-κB-driven responses (e.g., *FGFR2, RIPK3*, *NEK6*, *PRKGB*), highlighting distinct regulatory entry points into shared immune programs. *EGFR* and *ERBB2* (HER2) are canonical oncogenic drivers of MHC-I suppression^79,80^. Our results further align with clinical observations where *MAP3K1* deficiency impairs MHC-I-mediated antigen presentation in breast cancer^81^, while *ATR* inhibition has been shown to induce MHC-I expression and interferon signaling in lung cancer models^82^. Kinase perturbation analysis revealed that knockdown of *GSK3A*, *RIPK3, FGFR2* lead to a marked decrease in NF-κB expression, in agreement with prior studies positing these genes as positive regulators of NF-κB activity or GBM cell survival^83–85^. Additionally, IFN-γ signaling exhibited the greatest overlap in kinase regulators with the other pathways examined. This is consistent with the central role of IFN-γ in inducing both MHC-I and MHC-II expression^86^, while also engaging in crosstalk with NF-κB signaling^87^. Together, these results demonstrate that kinase perturbations exert modular yet convergent effects on tumor-immune interactions, with discrete nodes influencing MHC expression, interferon and NF-κB signalings (Supplementary Figure 4f).

**Figure 2.**
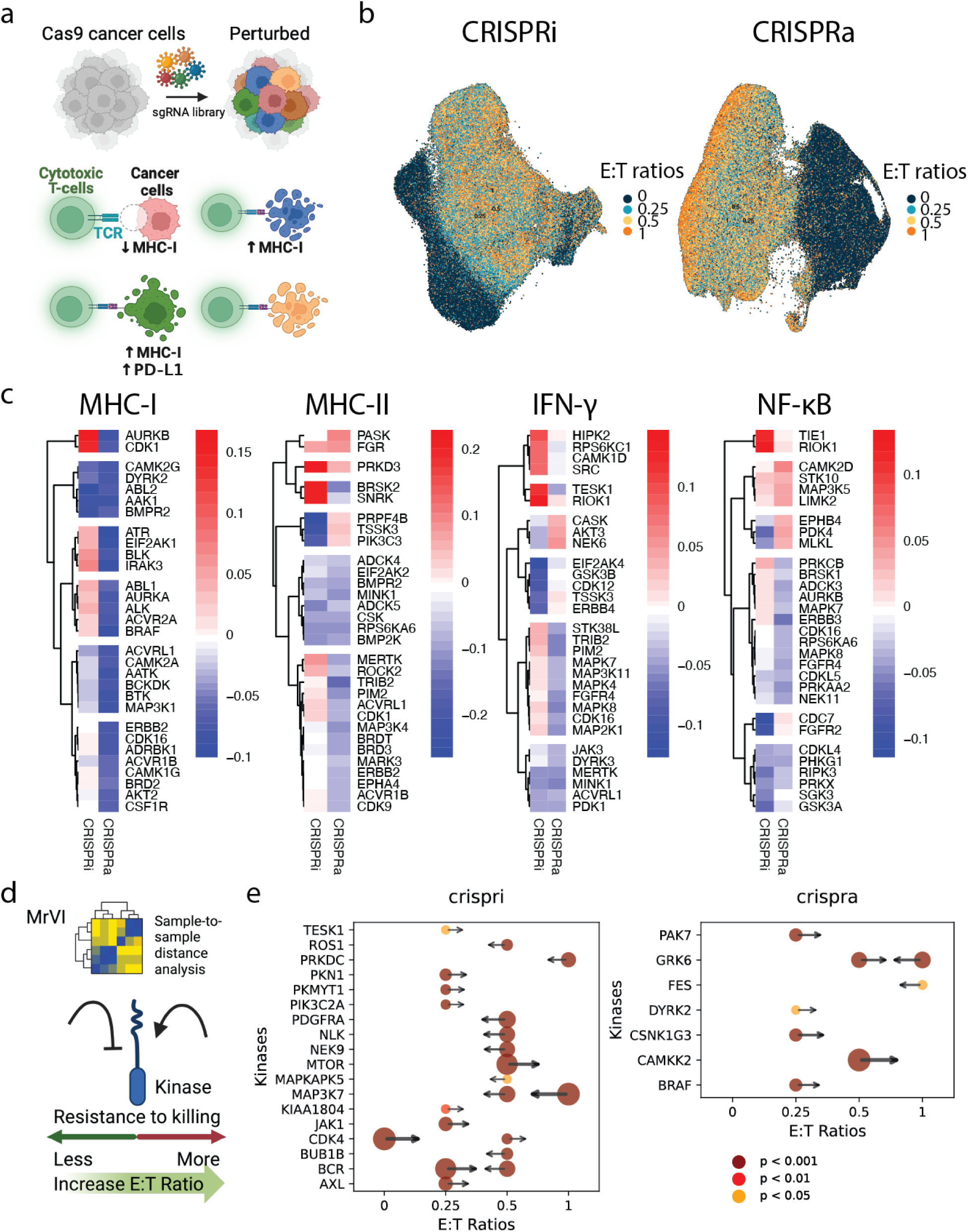
Kinase perturbations remodel antigen-presentation and immune response programs in GBM cells upon T cell exposure. (a) Experimental schematic of CRISPRi/a perturbations in NY-ESO-1/HLA-A*02:01-engineered GBM cells co-cultured with T cells. (b) UMAP embedding, generated from PCA space, of cancer cells under CRISPRi (left) or CRISPRa (right) showing transcriptomic trajectories across E:T ratios. (c) Module-level changes in MHC-I, MHC-II, IFN-γ, and NF-κB pathway programs under CRISPRi versus CRISPRa (top 30 kinases, FDR <0.05). Colors indicate effect size, quantified as the β coefficient for the genotype term (perturbation) in a regression model evaluated under the No T cell condition. (d) Schematic for MrVI analysis, and (e) Forest plots summarizing perturbation effects on the T cell–induced transcriptional program across effector:target (E:T) ratios, estimated by MrVI from global transcriptomic differences. Color and circle size denote p values, while arrow length and thickness represent distance differences. The statistical tests were performed with a one-sided Mann-Whitney U test (Bonferroni-corrected, α < 0.05).

We next used negative binomial regression to examine the effect of kinase perturbation on the wider T cell-induced program (Figure 1c). Our analysis identified 205 kinases (133 from CRISPRi and 93 from CRISPRa, 21 shared) whose perturbation significantly altered the top 50 genes of the program (see Methods, Supplementary Figure 4e,5f, Supplementary Table 4, 5). To more quantitatively assess the effects of kinase perturbation on modulating the global transcriptomic shifts induced at increasing E:T ratios, we used multi-resolution variational inference (MrVI)^88^ to define the ability of a genetic perturbation to dampen or exacerbate the changes induced by T cell exposure.

MrVI analysis showed that perturbed GBM cells without T cell exposure clustered tightly for both CRISPRi and CRISPRa conditions (Supplementary Figure 6a-b,7a-b). We next quantified how closely each perturbed sample aligned with its native E:T condition versus other E:T groups as a measure of how a genetic perturbation alters dose-response relationships. Specifically, for each sample, we calculated the transcriptomic similarity to every E:T ratio group (0, 0.25, 0.5, 1) and identified cases where the sample was similar to a different E:T ratio group than its own (see Methods, Figure 2d). Notably, knockdown of *PDGFRA*, the second most amplified receptor tyrosine kinase in GBM^87,89^, shifted tumor cells toward transcriptomic states characteristic of lower E:T ratios, suggesting reduced tumor evasion to T cell-mediated pressure (Figure 2e, Supplementary Table 10,11). Similarly, *JAK1* deficiency has been reported to confer resistance to T cell-mediated killing^90^, consistent with our CRISPRi findings showing that loss of *JAK1* promotes immune evasion signatures. Conversely, BRAF inhibition has been shown to enhance T cell-mediated cytotoxicity in melanoma^91^, supporting our observation that *BRAF* activation increases signatures associated with immune evasion. These analyses reveal that specific kinase perturbations can disrupt the expected transcriptomic trajectory of GBM cells under T cell pressure.

### Kinase Perturbations Reshape T Cell-Induced Trajectories

To determine whether T cell-induced immune evasion reflects discrete transcriptional states or a continuous, ordered tumor cell response, we applied Decipher^92^ to model tumor-intrinsic trajectories under increasing immune pressure. Decipher is a deep generative model that learns and aligns trajectories in an interpretable 2D space, capturing both shared and unique transcriptional programs while preserving the global geometry of T cell-induced trajectories. This framework enables the characterization of cancer cell response trajectories and provides a platform by which to align CRISPRi and CRISPRa trajectories in the same latent space.

Across unperturbed cells, Deciper revealed continuous progression of tumor cell states as T cell pressure increased (Figure 3a,b, Supplementary Figure 8a), recapitulating the ratio-dependent effects described above. Along the inferred trajectory of cancer cell responses to increasing T cell exposure, we observed a structured reorganization of tumor-intrinsic transcriptional programs (*CSF3*, *SOD2*, *IDO1*, *CD274*, *AR*, and *ID1,* Figure 3c, Supplementary Figure 8b). Additionally, genes significantly upregulated across the trajectory (Supplementary Table 12) included interferon-responsive innate immune signaling *(STAT1, IRF1, GBP1, GBP2*)^50,93^, antigen presentation machinery (*HLA-B, HLA-DRA, CIITA, TAP1, TAP2*)^79,94^, co-occurring with NF-κB signaling pathway (*NFKB1*, *NFKBIZ*, *TNIP1*)^59,95^ and chemokine signaling (*CXCL2, CXCL3, CXCL8, IL1B, IL6*)^96,97^. Additionally, we observed increased anti-apoptic, stress-resistance, and immune evasion associated genes like *SOD2*, *NAMPT*, *IDO1*, as well as the presence of ferroptosis regulators (*FTL^98^, SLC39A14^99^*). This aligns with the established role of IFN-γ-driven ferroptosis in T cell-mediated killing of GBM cells, suggesting that resistance to ferroptosis may represent an additional mechanism of GBM immune evasion^100^. Lastly, genes promoting stemness (*ID1*, *ID3*)^67^, proliferation (*MYLK*)^70^, and migration (*ENAH*)^101^ were downregulated along the trajectory (Figure 3c, Supplementary Table 12).

**Figure 3.**
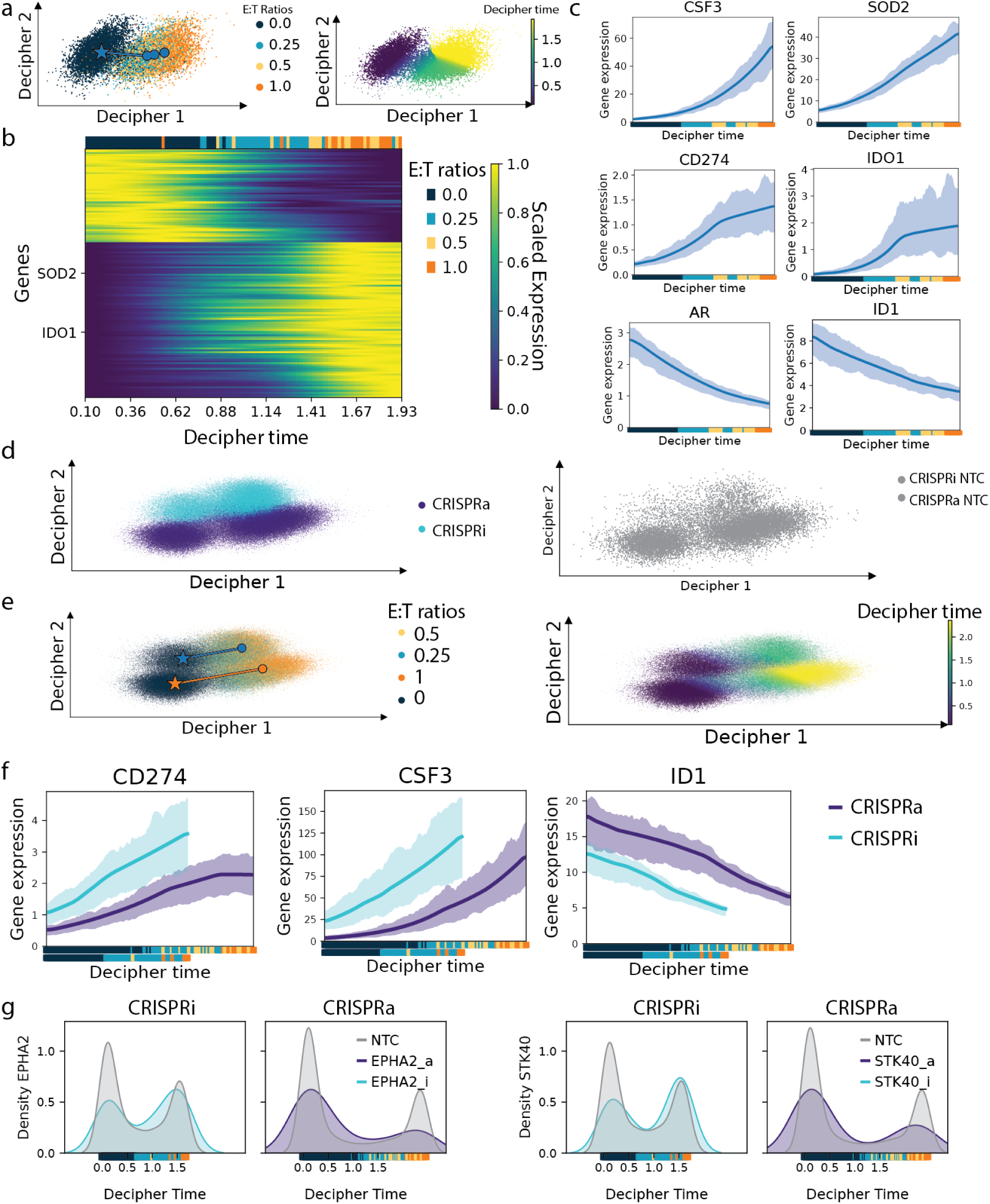
Decipher analysis of CRISPRi/a perturbations in GBM-T cell co-cultures reveals kinase dependent shifts in the T cell-induced program. (a) Decipher analysis of NTC cells from CRISPRa data reveals trajectories driven by increasing T cell concentration, shown in the learned 2D Decipher space with cells colored by E:T ratios (left) and corresponding Decipher time (pseudotime; right). (b) Genes significantly contribute to the Decipher trajectory, with the positions of *IDO1* and *SOD2* highlighted. (c) Single-cell trajectories of representative genes inferred by Decipher reveal E:T ratio-dependent transcriptomic responses. (d) Decipher-aligned CRISPRi and CRISPRa cells (left) projected into a shared 2D Decipher space together with NTC cells (right). (e) Decipher trajectories learned as a function of E:T ratio (left) and pseudotime (right) for CRISPRi and CRISPRa conditions. (f) Comparison of CRISPRi and CRISPRa perturbations highlights conserved effects on immune checkpoint expression (*CD274*) and tumor-intrinsic T cell-responsive genes (*CSF3*). (g) Density plots illustrate distribution shifts of cancer cells along Decipher time following perturbation of selected kinases (*EPHA2*, *STK40*), indicating that kinase disruption alters the progression of cancer cell states under T cell pressure (p = 0.033639216, p = 0.008503096 with Kolmogorov-Smirnov test).

Having established the ability of Decipher to recreate the biological changes induced by T cells, we next applied it to jointly align model trajectories from CRISPRi and CRISPRa perturbed cells alongside their respective NTC, revealing strong concordance across conditions (Figure 3d). In both CRISPRi and CRISPRa datasets, we observed ratio-dependent trajectories that closely resembled those of unperturbed NTC cells (Figure 3e). Representative gene signatures (*SOD2*, *CSF3*, *CD274*, *IDO1* and *IL6* Figure 3f, Supplementary Figure 9a) along these aligned trajectories further captured conserved tumor cell responses to increasing T cell pressure.

This framework enabled us to study how kinase perturbations influence cancer cell responses to increasing T cell exposure, by examining the distribution of cancer cells along Decipher time (Supplementary Table 13,14). Notably, knockdown of *EPHA2* (p=0.044) shifted the distribution toward higher E:T ratios, whereas their overexpression produced the opposite effect (Figure 3g, Supplementary Figure 9b). Ephrin–Eph receptor interactions caused by cell-cell contact regulate cell proliferation and migration during development and in tissue homeostasis. A similar pattern was observed for *STK40*. Both *EPHA2* and *STK40* are implicated in cancer progression and therapy resistance^102,103^. We also identified modulation of the T cell-induced trajectory by *IRAK1* and *IRAK4*, implicating IL-1R/TLR inflammatory signaling^104^, as well as *STK17B*, and *STK35*, consistent with engagement AKT signaling pathways and tumor progression^105,106^ (Supplementary Figure 9b).

Collectively, these results highlight that tumor-intrinsic kinases do not simply modulate static immune evasion phenotypes, but actively reshape the trajectory of cancer cell responses to T cell pressure. By shifting tumor cells toward states that either amplify or dampen immunogenic programs, these kinases represent potential levers for therapeutically sensitizing tumors to T cell-mediated killing for understanding mechanisms of immune resistance in GBM.

### Small molecules inhibition of select kinases modulate the T cell-induced program in patient-derived neurospheres

Because Decipher and MrVI increase power to detect kinases that shift the global T cell–induced transcriptional program, but do not directly specify which program subsets are affected, we coupled these analyses to an iterative compound-screening strategy to functionally resolve effects on immune-evasion programs and tumor susceptibility to T cell-mediated killing. Integrating Decipher with generalized linear modeling and MrVI, we prioritized kinases (*IRAK4*, *NTRK3*, *CDK4*, *TTK*, *GSK3A*, *PDGFRA*, *EPHA2*, *NEK11*, *CDK2*, *STK16*, *BMP2K*, Supplementary Table 4,5,10,11,13,14) that appear to modulate the tumor-intrinsic response to cytotoxic T cell exposure and identified 11 chemical inhibitors targeting these kinases (Table 1, Figure 4a). To determine whether chemical inhibition can modulate the T cell-induced program in patient-derived GBM neurospheres, we engineered them to overexpress the NY-ESO-1 antigen and the HLA:02*01 allele to enable antigen-specific T cell recognition (Supplementary Figure 1b). Neurospheres were treated with small molecule inhibitors (0.01 μM-10 μM) for 48 hours, after which drugs were removed and cells were co-cultured with T cells for 18 hours followed by sci-Plex multiplexing and single-cell RNA-seq. Similarly, T cell exposure induced a dominant transcriptomic shift in PDN cells (Figure 4b). Cluster 2 was enriched for T cell-treated samples, indicating a shared T cell-driven transcriptional state. Notably, treatment with CP-673451 (PDGFRA inhibitor), STK16-IN-1 (STK16 inhibitor), abemaciclib mesylate (CDK4 inhibitor), and ALW-II-41-27 (EphrinA2 inhibitor) resulted in a reduced proportion of cells in Cluster 2 compared to DMSO, whereas most other compounds increased the representation of this T cell-associated cluster (Figure 4c, Supplementary Figure 10a).

**Figure 4.**
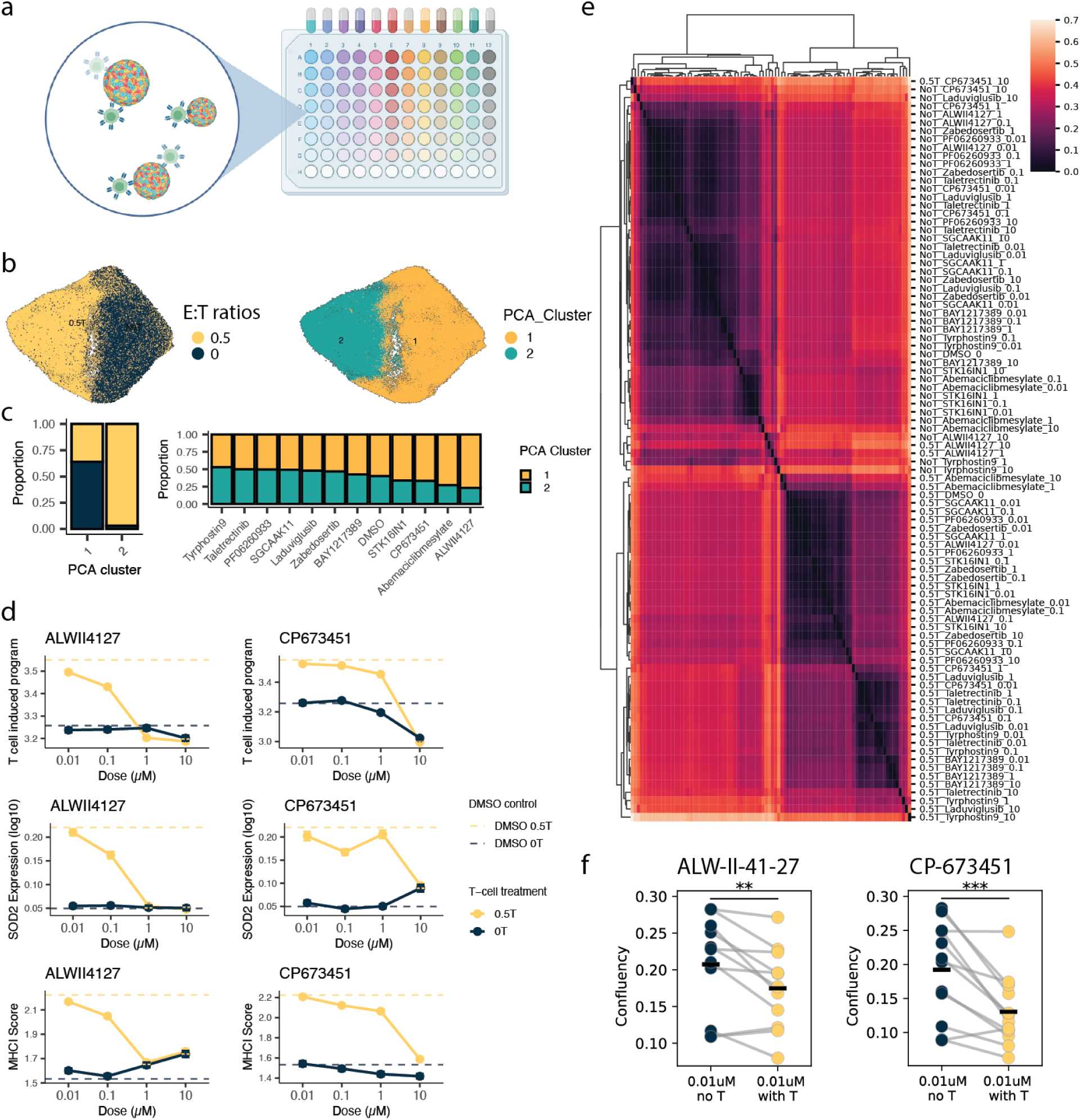
Chemical perturbation validation of immunomodulatory kinase targets in GBM-T cell co-cultures. (a) Schematic of chemical inhibitor screen design using GBM:T cell co-cultures. (b) UMAP visualization of the PCA space, colored by T cell treatment (E:T ratio = 0.5) vs No T cells (No T) (left) and PCA clusters (right). (c) PCA clustering of drug-treated samples, shown by E:T ratio distribution across clusters (left) and cluster assignment across different drugs (right). (d) Representative concentration-response curves for the PDGFRA and EPHA2 inhibitors CP-673451 and ALW-II-41-27, respectively, showing effects on T cell response score, *SOD2* expression, and MHC-I score in the presence or absence of T cells. (e) Sample-to-sample distance heatmap of drug perturbations, generated by MrVI, highlights distinct clustering of treatments. (f) Tumor cell confluency following treatment with CP-673451 or ALW-II-41-27 (0.01 µM) in the absence or presence of T cells. Paired measurements from the same experimental replicates are connected by gray lines; colored dots represent individual replicates, and horizontal bars indicate mean values. Cell confluency is expressed as a fraction of the well area covered by cells (e.g., 0.20 = 20% of the well is occupied by GBM cells). Statistical significance was assessed using paired t-tests (p values = 0.005, 0.0006 respectively).

**Table 1.**
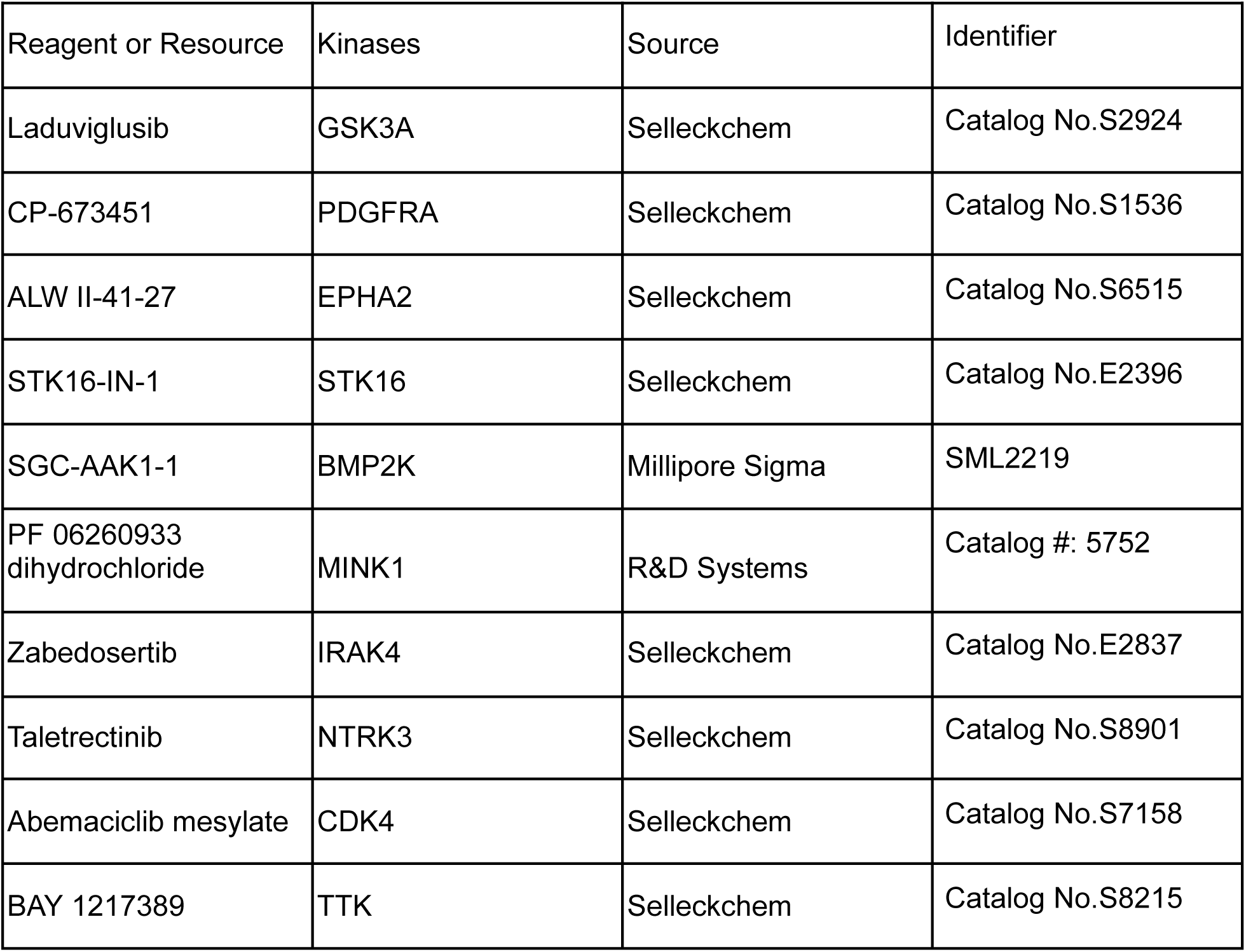
Kinase inhibitors.

Across our drug panel, we identified two drugs that with or without T cell treatment showed a major impact on PDN transcriptomics profiles at high concentration (1 μM and/or 10 μM). CP-673451 and ALW-II-41-27 exhibited strong attenuation of the T cell-induced programs, *SOD2*, *CD274*, and MHC-I and MHC-II programs (Figure 4d, Supplementary Figure 10b), an effect not observed with the other drugs tested (Supplementary Figure 11). Using MrVI to quantify pairwise transcriptomic distances across our treatments revealed that the highest concentration of these two drugs has the closest transcriptomic distance compared to No T cell controls (Figure 4e), consistent with our prior genetic results. Additionally, the MrVI latent (z) space was dominated by the effect of T cell exposure, with most drug-treated samples well mixed across conditions. In contrast, a subset of samples treated with CP-673451 and ALW-II-41-27 grouped with the No T cell treatment condition, indicating that these perturbations partially revert or attenuate the T cell-induced transcriptional state. (Supplementary Figure 12).

We next tested whether this shift in transcriptional program led to changes in the ability of cytotoxic T cells to target tumor cells for destruction using a T cell cytotoxicity assay in 2D patient-derived GBM cultures. Cells were treated with both compounds at 0.01 or 0.1 μM. We observed that low-dose treatment (0.01 μM) for ALW-II-41-27 and or both doses for CP-673451 significantly enhanced T cell-mediated killing, supporting our findings in 3D neurosphere cultures (Figure 4f, Supplementary Figure 13).

Lastly, we explored whether there was an association with PDGFRA copy number and the levels of immune evasion genes induced by T cell exposure. Analysis of published GLASS bulk RNA-seq data⁷⁵ revealed that PDGFRA copy number loss was associated with reduced expression of *IDO1*, *SOD2* and *CD274* (Supplementary Figure 14). Given the low number of samples with low PDGFRA copy number we are not able to investigate a similar association for the IDH-wild type glioblastoma subset alone. However, low grade gliomas are also considered immunogenically cold tumors and PDGFRA targeting may aid in blocking the expression of immune evasion genes widely across gliomas^107^. Notably, *PDGFRA* amplification coincided with elevated IDO1 and SOD2 levels (Supplementary Figure 15) in glioma samples from in the TCGA Pan-Cancer Atlas^108^. Consistent with our perturbation results, PDGFRA inhibition can potentially reduce tumor-intrinsic resistance mechanisms and enhanced susceptibility to T cell-mediated killing^109,110^. These results indicate that inhibition of PDGFRA and EphrinA2 signaling using CP-673451 and ALW-II-41-27, respectively, has the potential to enhance T cell-based immunotherapies in combination treatments.

## Discussion

This study establishes a scalable single-cell perturbation framework for dissecting tumor-immune crosstalk under graded cytotoxic T cell pressure. By integrating pooled CRISPRi/a perturbations with sci-Plex-based single-cell profiling and deep generative modeling, we provide a resource for systematically mapping how genetic perturbations modulate tumor-intrinsic immune responses, enabling the discovery of regulators that reprogram the dynamic trajectory of immune evasion rather than static endpoints of resistance.

We show that immune evasion unfolds as a continuous transcriptional process where the progression and direction can be quantitatively modulated by defined genetic perturbations, consistent with prior work demonstrating that GBM malignant cells occupy plastic transcriptional states rather than fixed identities^111^. Additionally, immune cells have been shown to adopt continuous transcriptional trajectories across numerous cancer types, including GBM^112,113^. Trajectory analyses using Decipher and MrVI revealed that tumor cells progress along a continuum of immune adaptation to stress-resilient and immune-evasive phenotypes. Within this landscape, our kinome-scale perturbation identified signaling nodes that re-route state progression. Among these kinases, *PDGFRA* and *EPHA2* emerged as key trajectory regulators that steer GBM cells away from immune-evasive paths. Perturbation of these kinases altered the pseudotemporal alignment of tumor cell responses to T cell pressure, consistent with tunable control points that can attenuate or exacerbate entry into immune-evasive programs depending on pathway context.

GBM cells exposed to T cells *in vitro* are efficiently killed, with strong cytokine responses^114^. *In vivo*, T cell therapies can suppress tumor growth and extend survival, but their efficacy is often limited by immune evasion mechanisms^115^. Overcoming these barriers is key to improving T cell-based therapies for GBM. The first clinical trial of EGFR-targeting CAR T cells in glioblastoma demonstrated both the safety of this approach and robust T cell infiltration within tumor tissue, yet also revealed a compensatory upregulation of immune-suppressive molecules, including IDO1 and PD-L1, consistent with adaptive resistance to T cell-mediated killing^32^. Additionally, Zhao et al., showed that responders to anti-PD-1 therapy in glioblastoma exhibited a distinct immune composition, transcriptional state, and clonal dynamics^26^. IDO1 is known to increase in GBM tumors due to T cell infiltration^62^. Additionally, upregulation PD-L1 seems to be associated with more aggressive GBM and correlated with T cell activation pathways in clinical samples^17^, which aligns with our findings in upregulation of *CD274* upon T cell exposure. Our kinome-scale perturbation screen addresses this gap by revealing specific kinases that govern how GBM cells sense and respond to T cell-derived immune pressure. These kinases (*NLK, CDK2*, *CAMK1D,* Supplementary table 4,5) act upstream of canonical immune-evasion programs, including PD-L1 and IDO1, providing actionable entry points to reprogram tumor susceptibility to T cell attack.

EPHA2 and PDGFRA are appealing translational targets in GBM because they are recurrently implicated in disease biology and are pharmacologically targetable. EPHA2, a receptor tyrosine kinase, is frequently overexpressed in various cancers and acts as a stress antigen recognized by certain T cell receptors, especially γδ T cells^116^. Studies have shown that Epha2 deletion in mice reversed T cell exclusion and sensitized tumors to immunotherapy, which is mediated through EPHA2/TGF-β/SMAD axis-dependent activation of prostaglandin endoperoxide synthase 2^117^. In GBM, the transmembrane receptor EPHA2 is frequently upregulated and has been associated with poor prognosis and reduced overall survival^118,119,120^. EPHA2 has been targeted in cancer via several strategies, including small molecule kinase inhibitors, monoclonal antibodies, antibody-drug conjugates, and immunotherapies such as CAR-T or CAR-NK cells engineered to recognize EPHA2 on tumor cells^121–125^. CAR-T and CAR-NK cell therapies leverage the immune system to selectively kill EPHA2-expressing tumor cells, showing promising results in preclinical models^121^. Small molecule inhibitors (including ALW-II-41-27) block EPHA2’s kinase activity and can overcome drug resistance or enhance the effects of other therapies^126^. Thus, combination therapies, such as pairing EPHA2 inhibitors with other targeted agents (e.g., KRAS or HDAC inhibitors), can further enhance anti-tumor efficacy and overcome resistance mechanisms^127–129^.

PDGFRA has also been a prominent therapeutic target in glioblastoma. Multiple studies have explored both small molecule inhibitors and antibody-based strategies to inhibit PDGFRA signaling in GBM^130,131^. Avapritinib, a highly selective and brain-penetrant PDGFRA inhibitor, has shown clinical responses in patients with PDGFRA-altered high grade glioma, including partial and near-complete tumor regression in some cases, and has demonstrated good central nervous system penetration and tolerability^109^. Consistent with findings from both GLASS and TCGA, inhibition of PDGFRA could reduce resistance to T cell-mediated killing. However, PDGFRA amplification is a hallmark of the proneural GBM state^111^ and generally anti-correlates with mesenchymal inflammatory programs^132^, which are enriched for immune-interacting and PD-L1–high phenotypes^133^. As a result, subtype composition may confound bulk RNA-seq based associations between PDGFRA signaling and immune evasion.

Notably, Gai et al. discovered that co-upregulation of PDGFRA and EPHA2 led to worse patient prognosis and poorer therapeutic effects in GBM. Due to PDGFRA-induced EPHA2 activation, EPHA2 driven resistance mechanisms can limit the effectiveness of PDGFRA-targeted monotherapy^134^, prompting research into combination strategies. This finding further supports the identification of both targets. Overall, while targeting PDGFRA and EPHA2 in GBM has shown promise, especially in genetically defined subgroups, overcoming resistance and improving patient outcomes remain active areas of investigation.

Our study interrogates a well-controlled, simplified axis of tumor-immune interactions, and antigen-specific cytotoxic T cell pressure on engineered GBM models under short-term co-culture. This provides a reductionist system intentionally designed to enable scalable, high-throughput perturbation mapping under graded immune pressure. Looking forward, a major opportunity is to incorporate more complex tumor microenvironment niches^135–137^ *in vitro* in a manner that remains compatible with pooled perturbation screens and massively multiplexed single-cell readouts. The integration of such platforms could further delineate more complex context-specific vulnerabilities towards nominating novel strategies that potentiate immunotherapy by preventing (or rerouting) the emergence of immune-evasive trajectories. Together, our results delineate a continuum model of tumor immune adaptation governed by kinase signaling, demonstrating how tumor-intrinsic kinase perturbations reshape tumor–immune interactions and bridge oncogenic pathways with immune evasion dynamics. Beyond the specific findings in glioblastoma, the experimental and analytical framework presented here serves as a generalizable resource for decoding tumor-immune interactions across cancer types and for designing combination therapies that reprogram immune trajectories.

## Supplementary figures

**Supplementary Figure 1.**
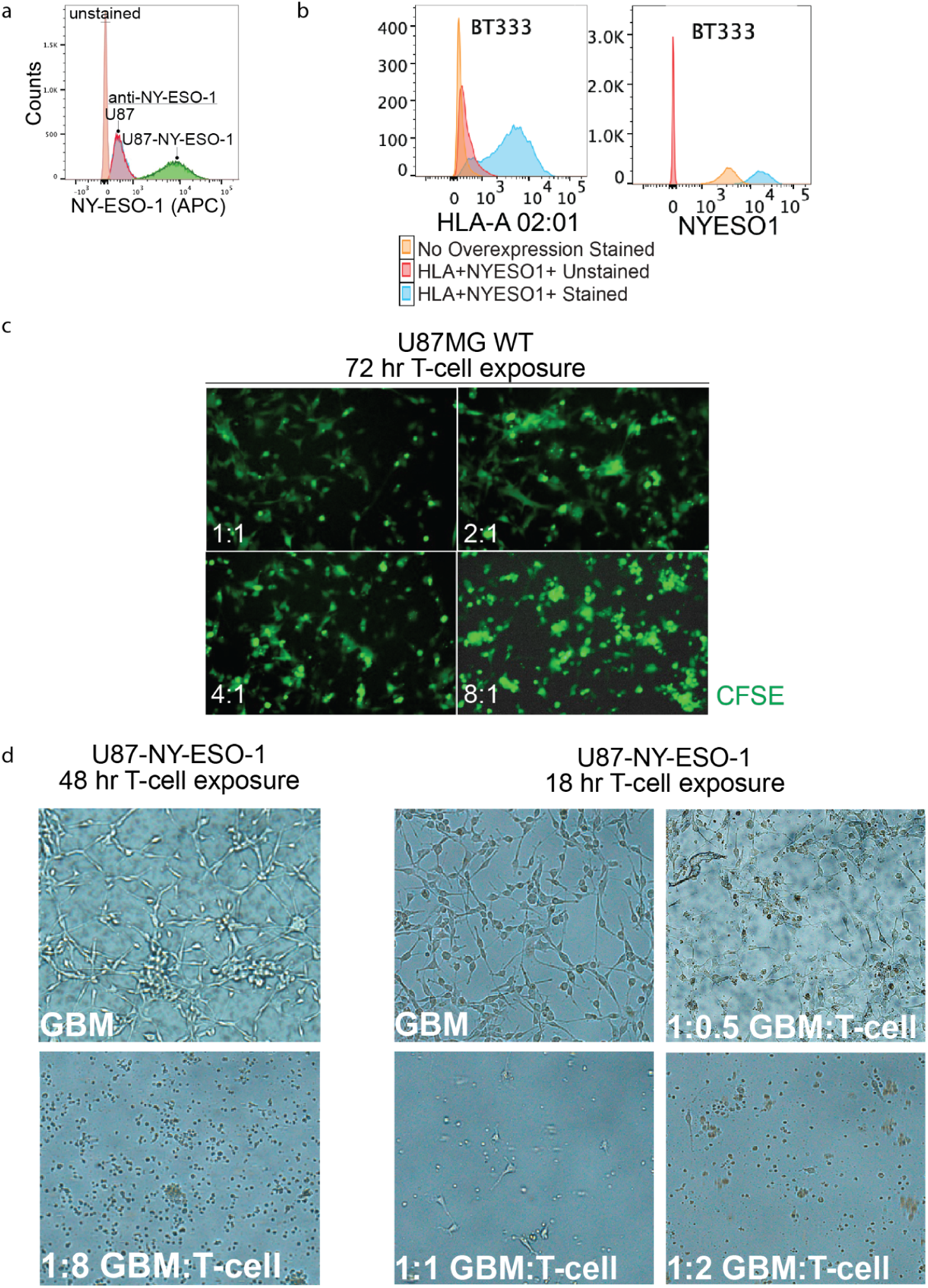
Engineering and Functional Validation of NY-ESO-1–Positive, HLA-Matched GBM Cells for T cell Co-culture. (a) Flow cytometry of NY-ESO-1 protein levels in U87MG cells with and without NY-ESO-1 overexpression. (b) Overexpression verification of HLA-0201 and NY-ESO-1 in the patient-derived GBM cell line BT333. (c) Fluorescent images of T cell:GBM co-cultures after 72 hours (CFSE, green). (d) Images of GBM NY-ESO-1/NY-ESO-1 TCR cytotoxic T cell co-cultures after 48 hours (left) and 18 hours of co-culture (right) at varying effector:target ratios.

**Supplementary Figure 2.**
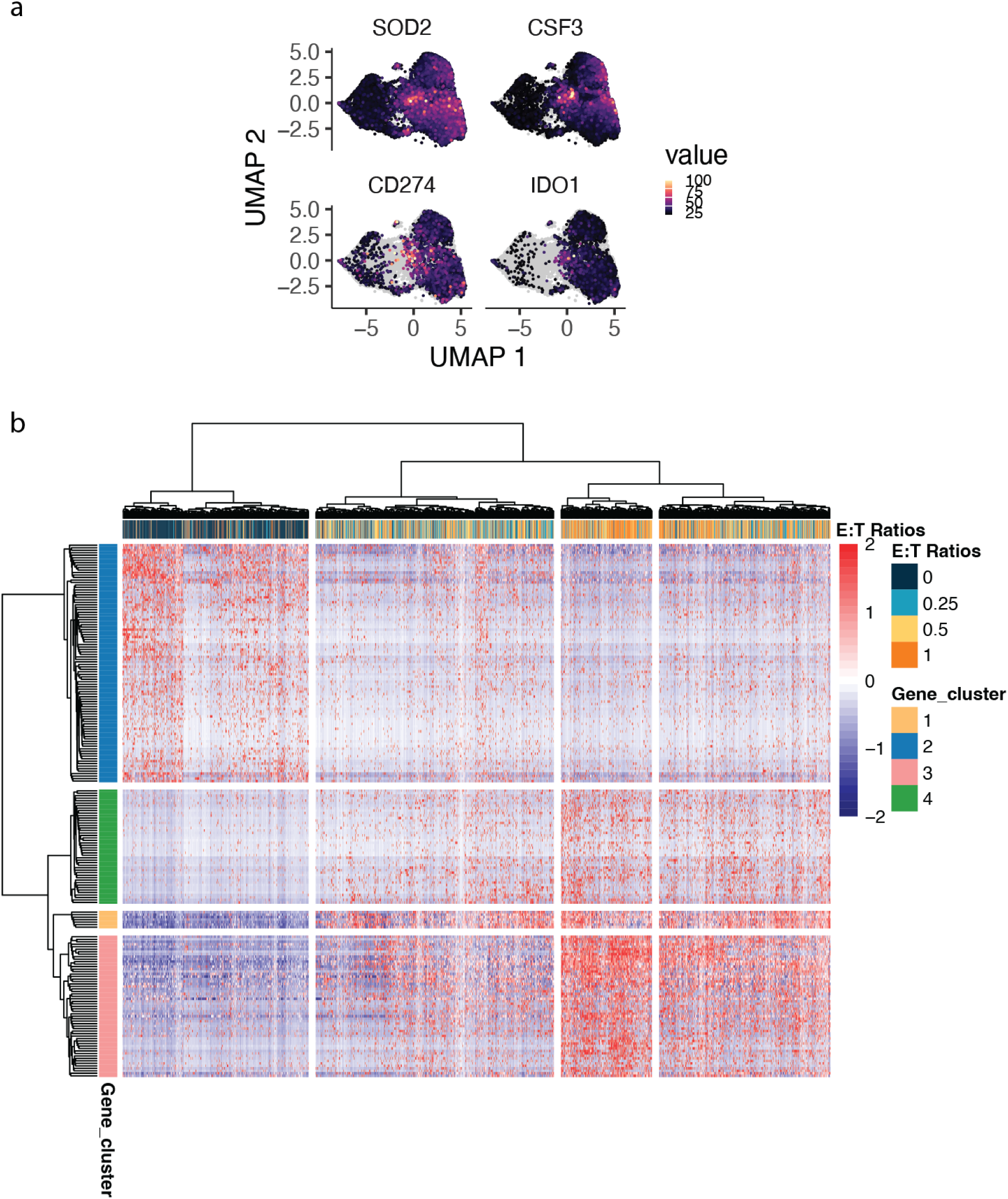
Differential expression analysis of unperturbed tumor cells reveals T cell induced programs. (a) UMAP visualization of the PCA space of *SOD2*, *CSF3*, *CD274*, and *IDO1* expression in non-targeting control cells. (b) Heatmap showing a graded T cell-induced transcriptional program across cancer cells.

**Supplementary Figure 3.**
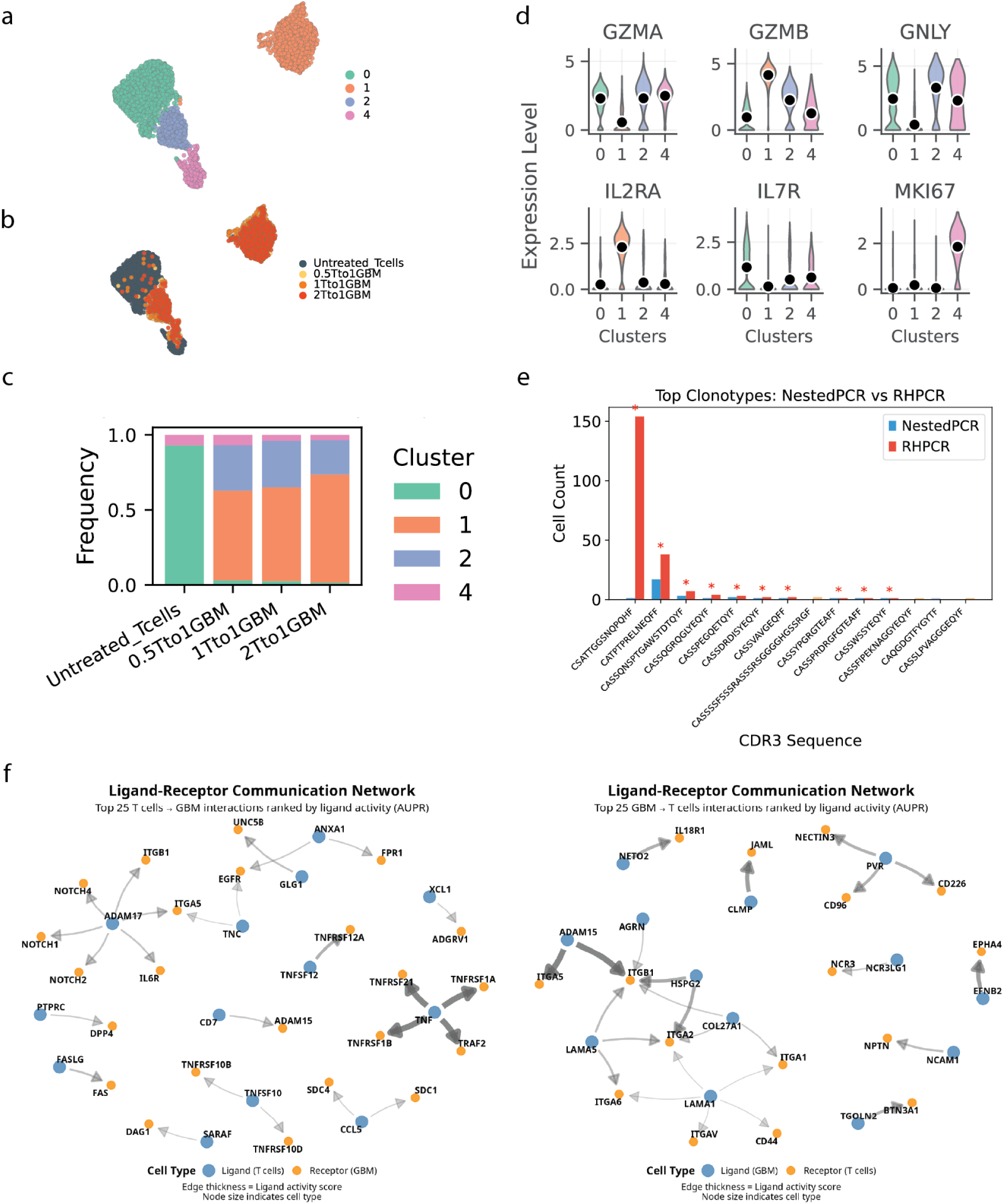
T cells responses upon exposure to GBM cells. (a, b) UMAP visualization of the PCA space of T cells from different E:T ratios, colored by (a) cluster or (b) E:T ratios. (c) Stacked bar plot showing cluster proportion for each E:T ratio. (d) Violin plots showing representative cytotoxic T cell signature gene expression across clusters. (e) TCR clonotype composition from two PCR enrichment approaches (Nested PCR, and RHPCR) (f) T cell to GBM and GBM to T cell ligand and receptor communication network (* means the clone existed in both TCR enrichment approaches).

**Supplementary Figure 4.**
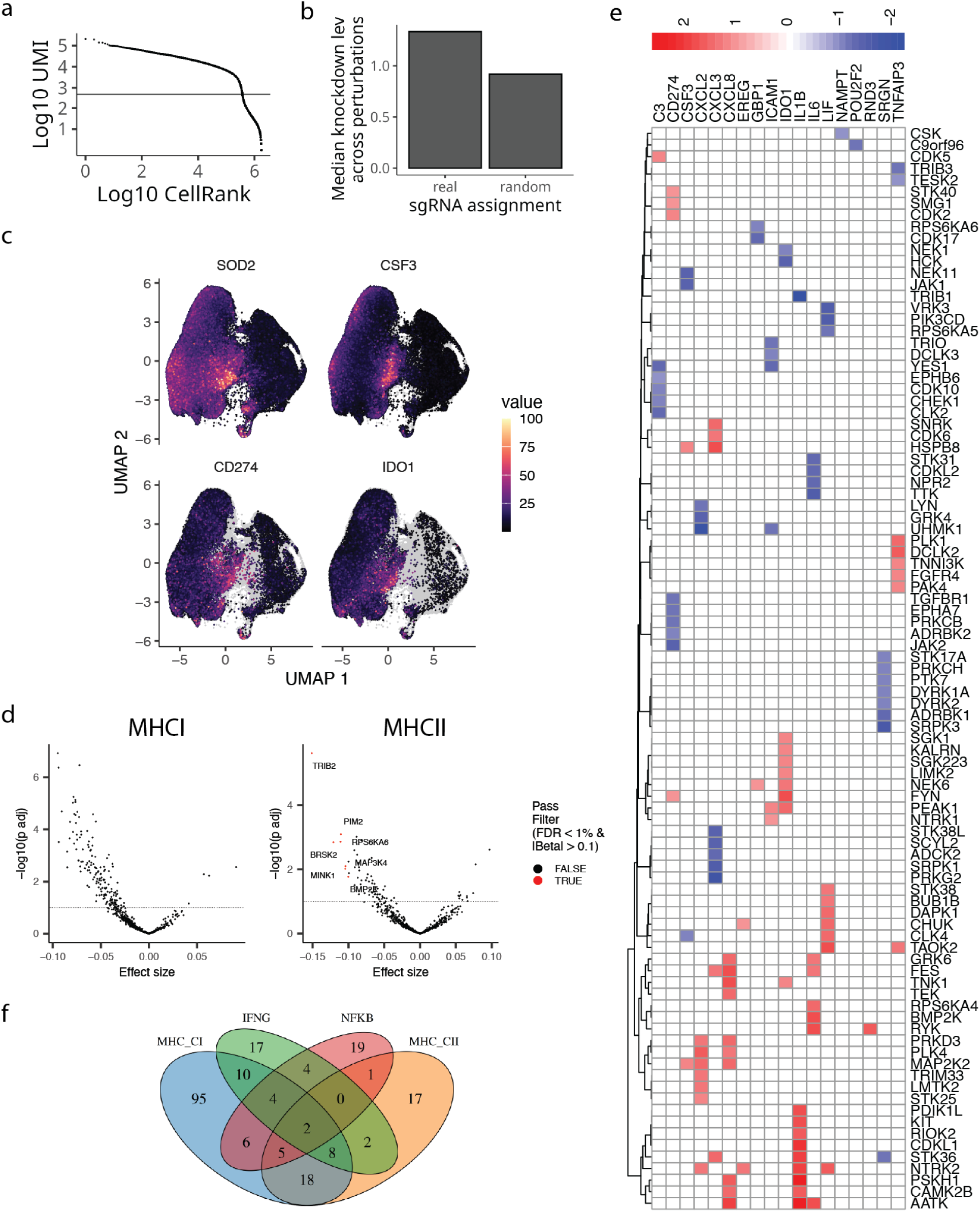
CRISPRa screen targeting kinases modulates tumor-intrinsic transcriptional responses in GBM cells. (a) Knee plot for CRISPRa data. (b) Median knockdown effect across sgRNAs targeting kinases of interest compared with random sgRNA assignment. (c) UMAP visualization of the PCA space of GBM cells colored by expression of representative T cell-induced program genes(*SOD2*, *CSF3*, *CD274*, and *IDO1*). (d) Volcano plots showing differential kinase regulation of MHC class I (left) and MHC class II (right) antigen presentation genes in response to T cell co-culture; highlighted genes pass significance thresholds (FDR < 1% and |β| > 0.1) (Supplementary Table 6,7). (e) Heatmap summarizing kinase perturbations that modulate T cell-induced programs. Colors indicate direction and magnitude of transcriptional effects, with hierarchical clustering. (f) Venn diagram comparing significant genes from Figure 2c (kinases that regulate MHC-I, MHC-II, IFN-γ, and NF-κB pathways).

**Supplementary Figure 5.**
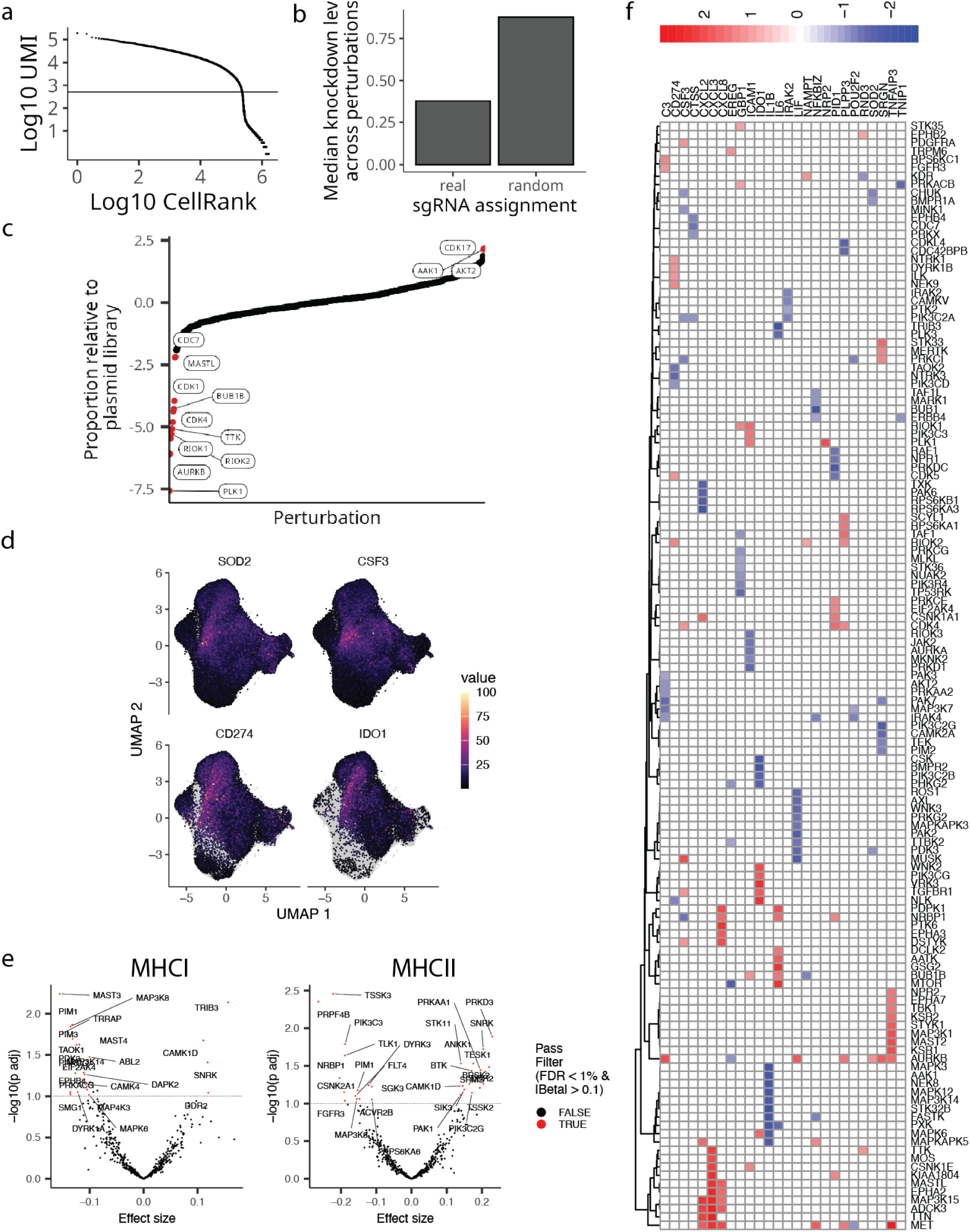
CRISPRi screen targeting kinases modulates tumor-intrinsic transcriptional responses in GBM cells. (a) Knee plot for CRISPRi data. (b) Median knockdown effect across sgRNAs targeting kinases of interest compared with random sgRNA assignment. (c) Guide proportion relative to plasma library; highlighted guides have log(relative proportion) > 2. (d) UMAP visualization of the PCA space of GBM cells colored by expression of representative T cell-induced program genes (*SOD2*, *CSF3*, *CD274*, and *IDO1*). (e) Volcano plots showing differential kinase regulation of MHC class I (left) and MHC class II (right) antigen presentation genes in response to T cell co-culture; highlighted genes pass significance thresholds (FDR < 1% and |β| > 0.1) (Supplementary Table 8,9). (f) Heatmap summarizing kinase perturbations that modulate T cell-induced programs. Colors indicate direction and magnitude of transcriptional effects, with hierarchical clustering.

**Supplementary Figure 6.**
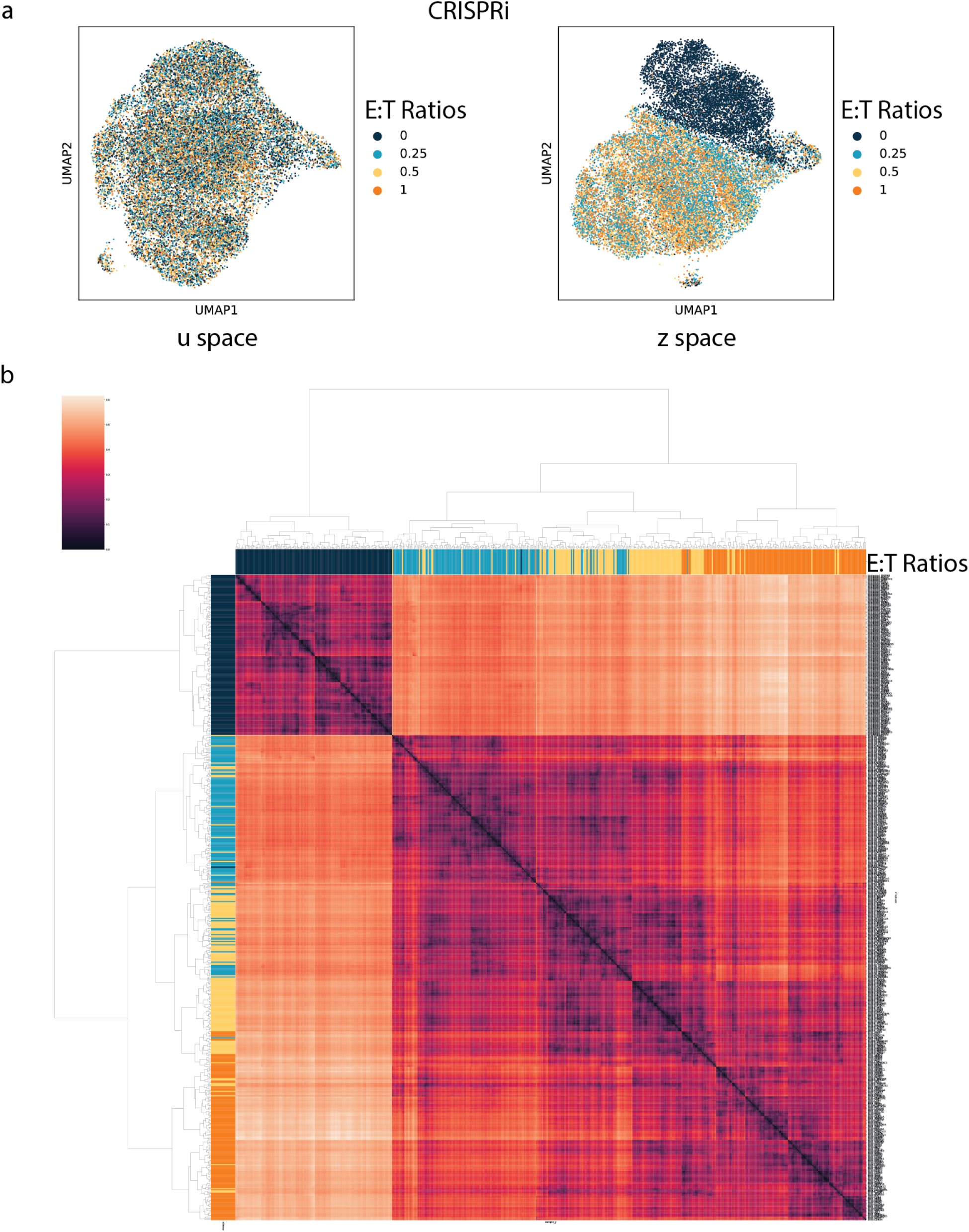
MrVI analysis of the CRISPRi kinome screen. (a) UMAP visualization of GBM cells under CRISPRi kinome screen in (left) u-space and (right) z-space, colored by E:T ratios. (b) Heatmap showing grouping of samples (E:T ratio and perturbation) based on MrVI-derived global transcriptomics distances.

**Supplementary Figure 7.**
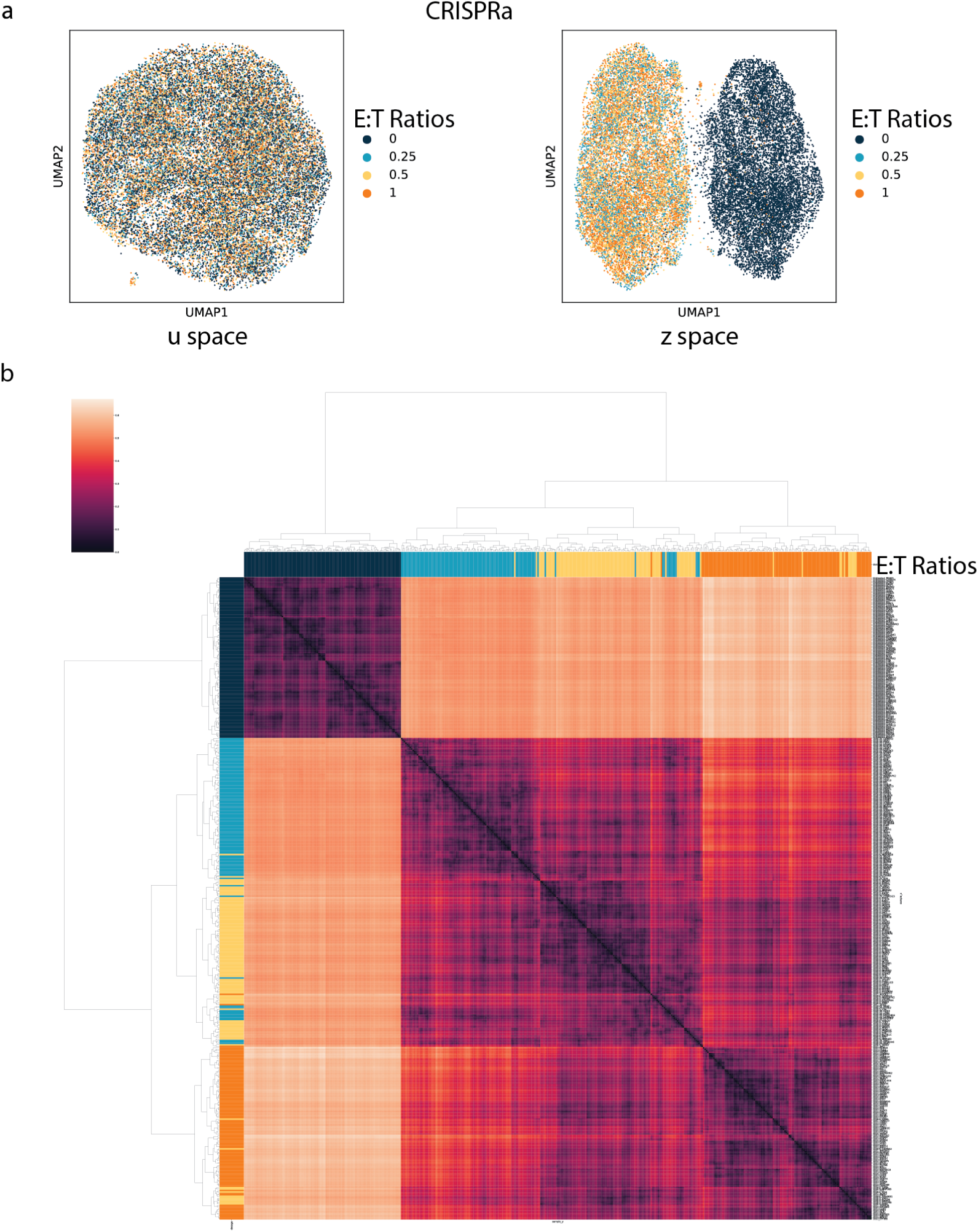
MrVI analysis of the CRISPRa kinome screen. (a) UMAP visualization of GBM cells under CRISPRa kinome screen in (left) u-space and (right) z-space, colored by E:T ratios. (b) Heatmap showing grouping of samples (E:T ratio and perturbation) based on MrVI-derived global transcriptomics distances.

**Supplementary Figure 8.**
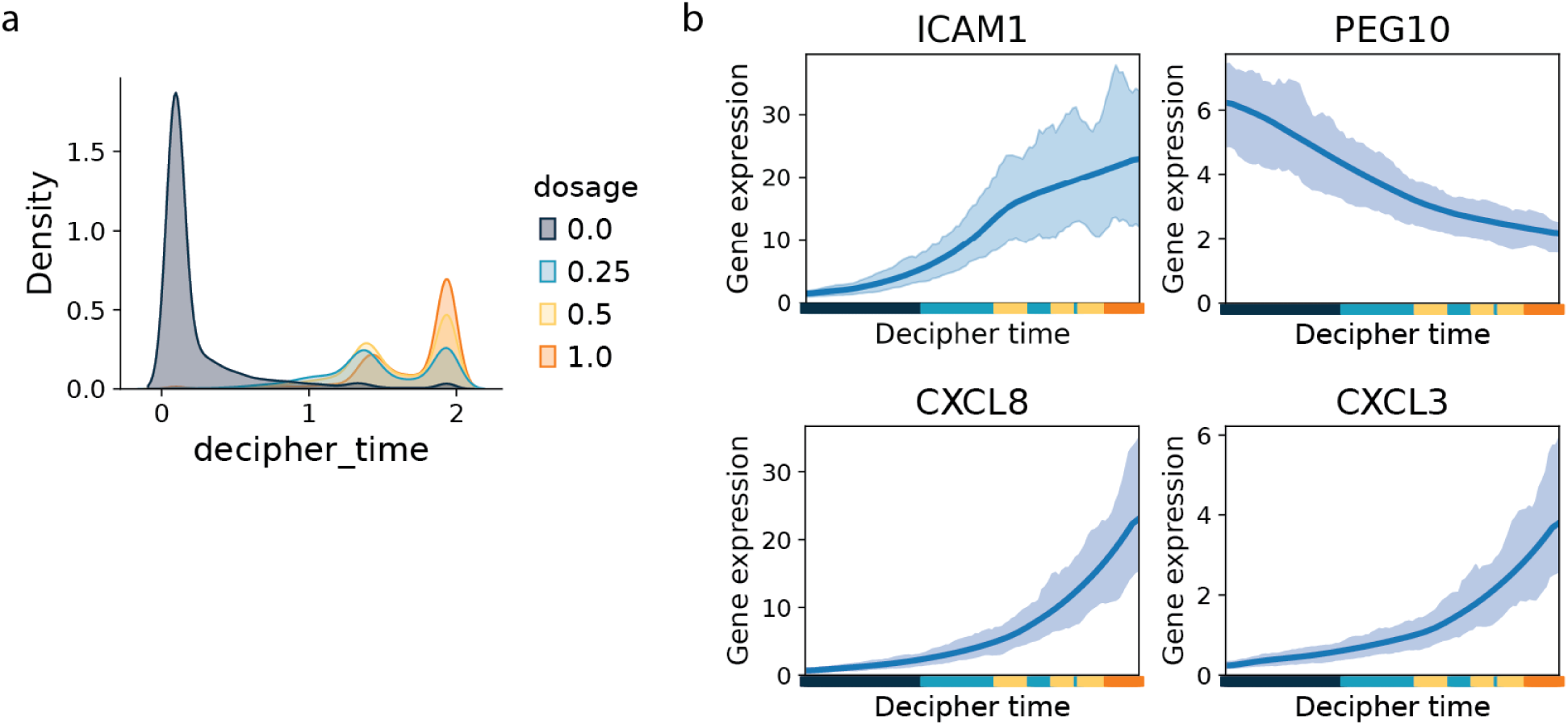
Decipher analysis of non-targeting controls reveals a T cell-induced cancer cell immune evasion program. (a) Density plot showing the distribution of E:T ratios along the Decipher trajectory. (b) Expression of *ICAM1*, *PEG10*, *CXCL8*, and *CXCL3* along the Decipher trajectory.

**Supplementary Figure 9.**
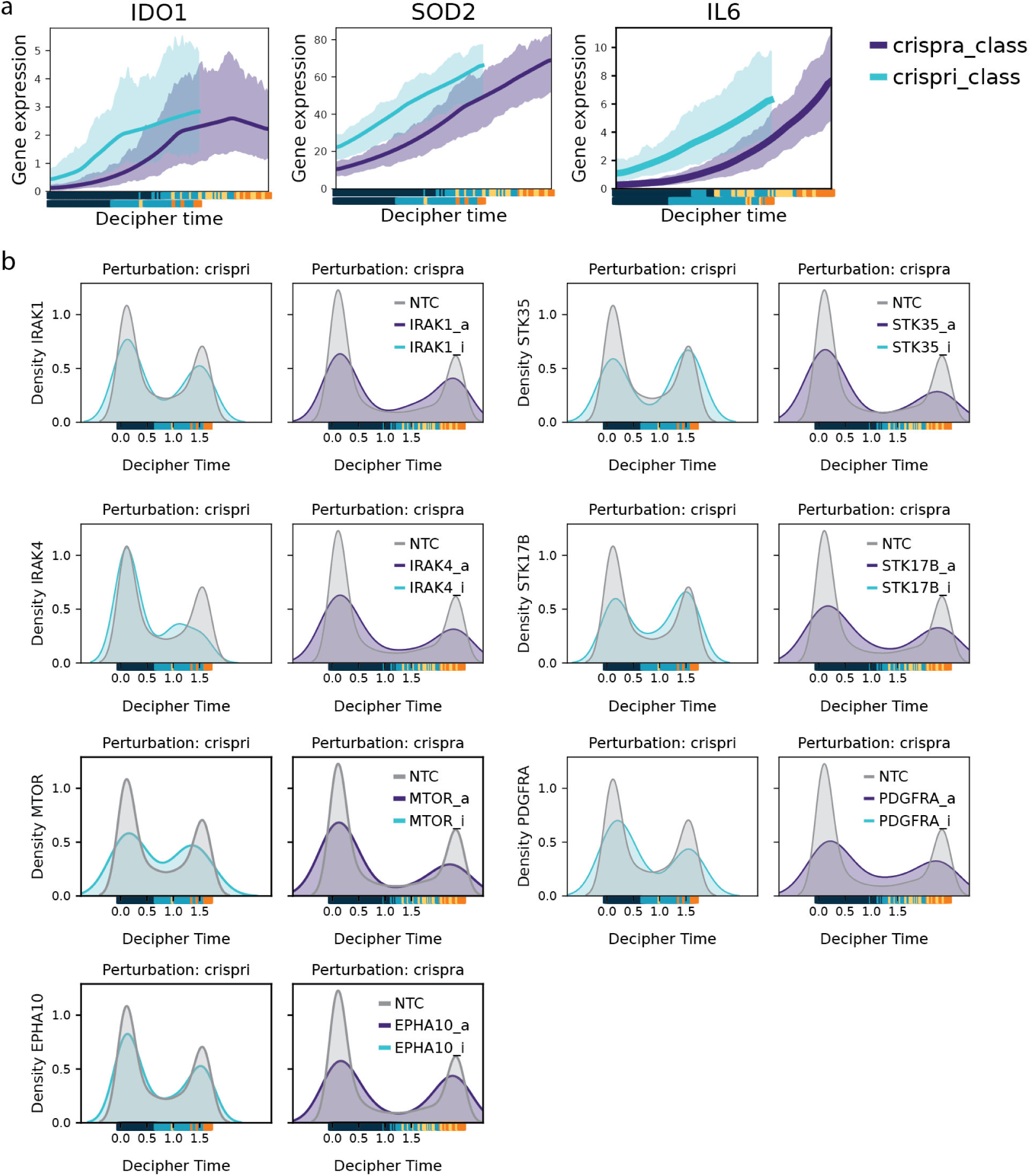
Decipher analysis of kinome perturbed CRISPRi & CRISPRa cells defines how perturbation affects the T cell-induced program. (a) Expression of *IDO1*, *SO2*, and *IL6* along the Decipher trajectory. (b) Density plots showing the distribution shift along the Decipher trajectory induced by perturbations of kinases (Supplementary 13,14).

**Supplementary Figure 10.**
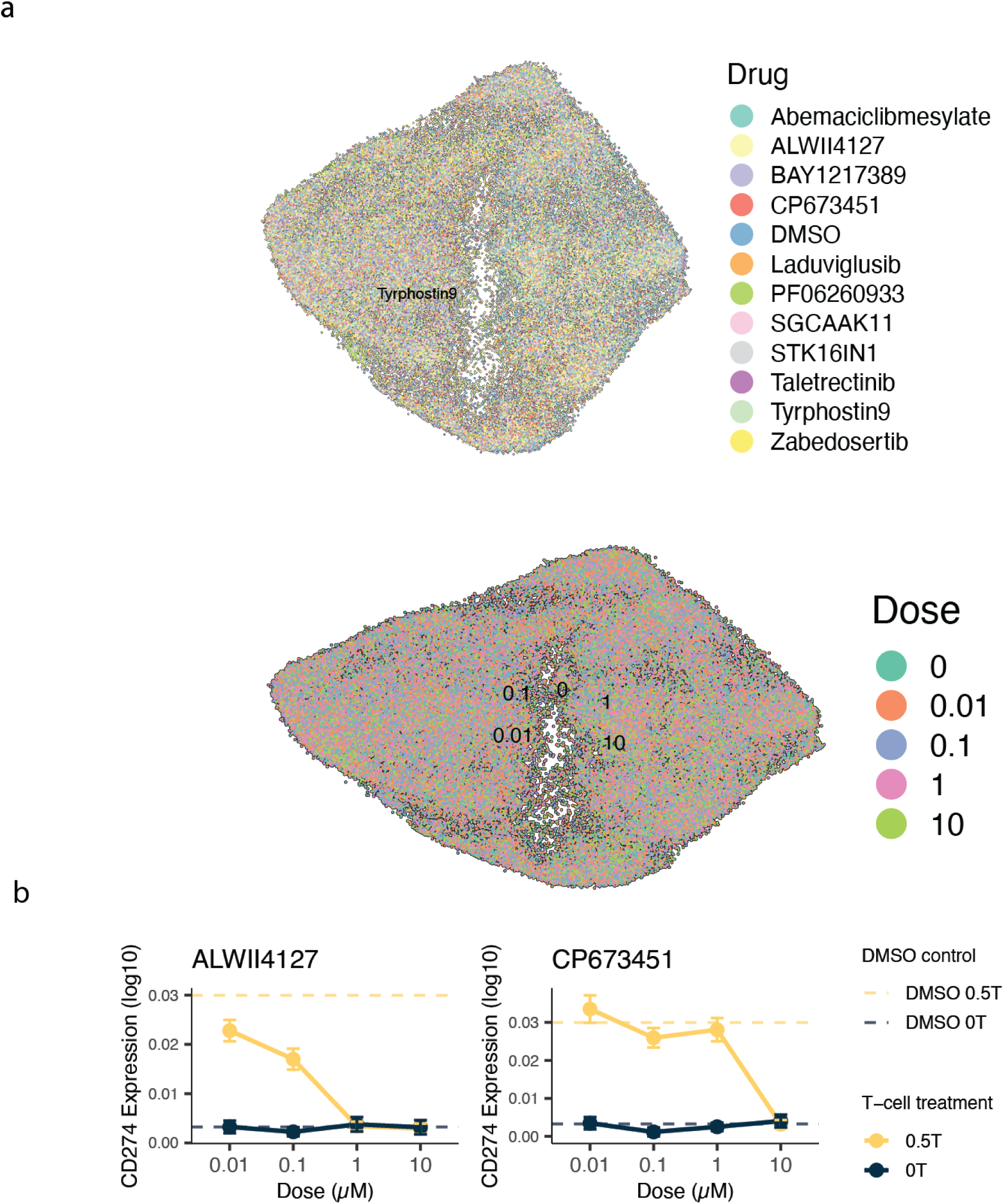
Drug- and dose-dependent effects in patient-derived neurosphere cells. (a) UMAP visualization of the PCA space of patient-derived neurosphere (PDN) cells colored by drug treatment (top) and dose (bottom). Cells that are 0 dose correspond to DMSO controls. (b) Dose-response curves for EphA2 and PDGFRA inhibitors showing effects on T cell *CD274* expression in the presence or absence of T cells.

**Supplementary Figure 11.**
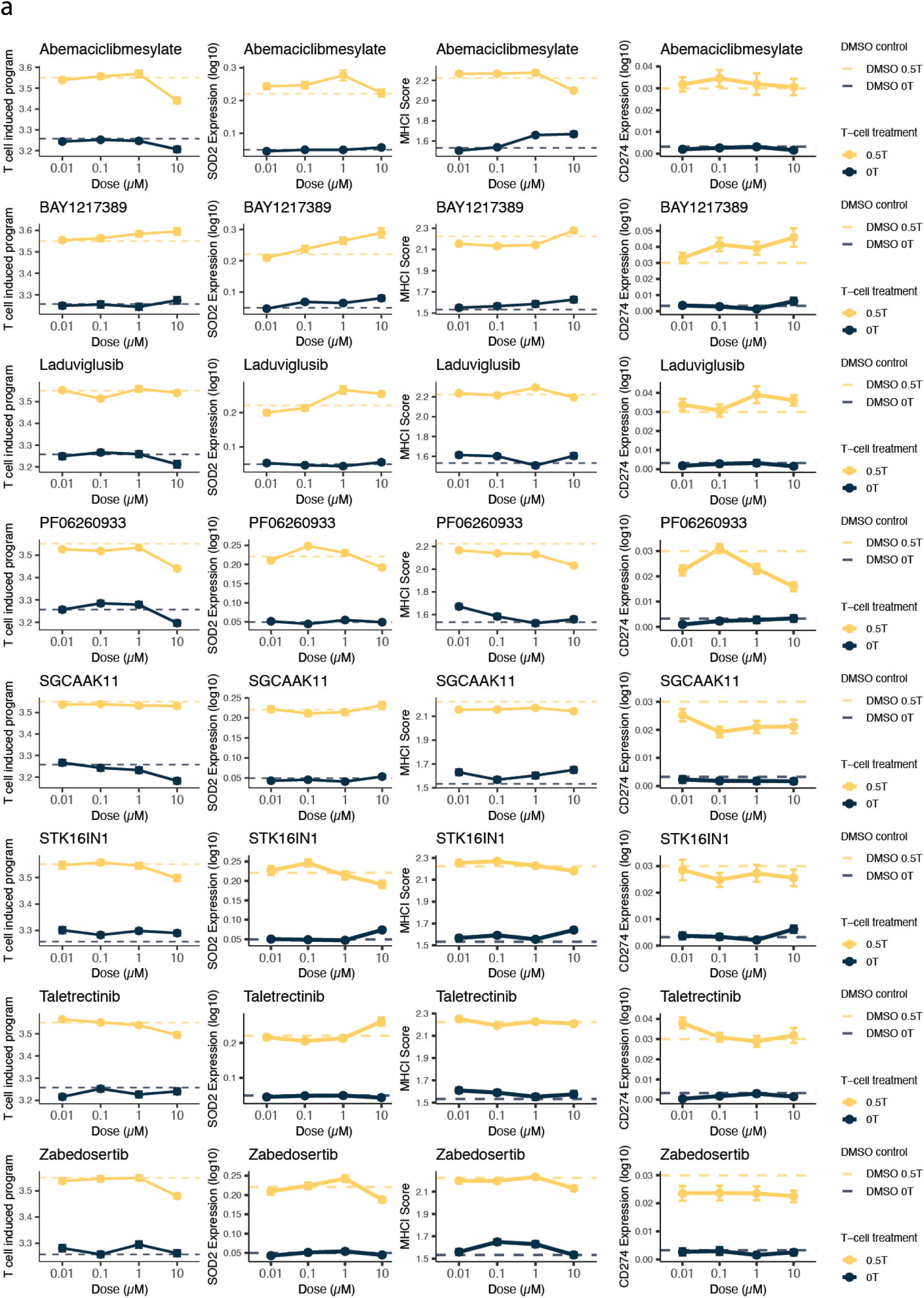
Dose-dependent effects of kinase inhibitors on T cell–induced programs in PDN cells. Representative dose–response curves for inhibitors showing effects on T cell response score, *SOD2* expression, and MHC-I score in the presence or absence of T cells.

**Supplementary Figure 12.**
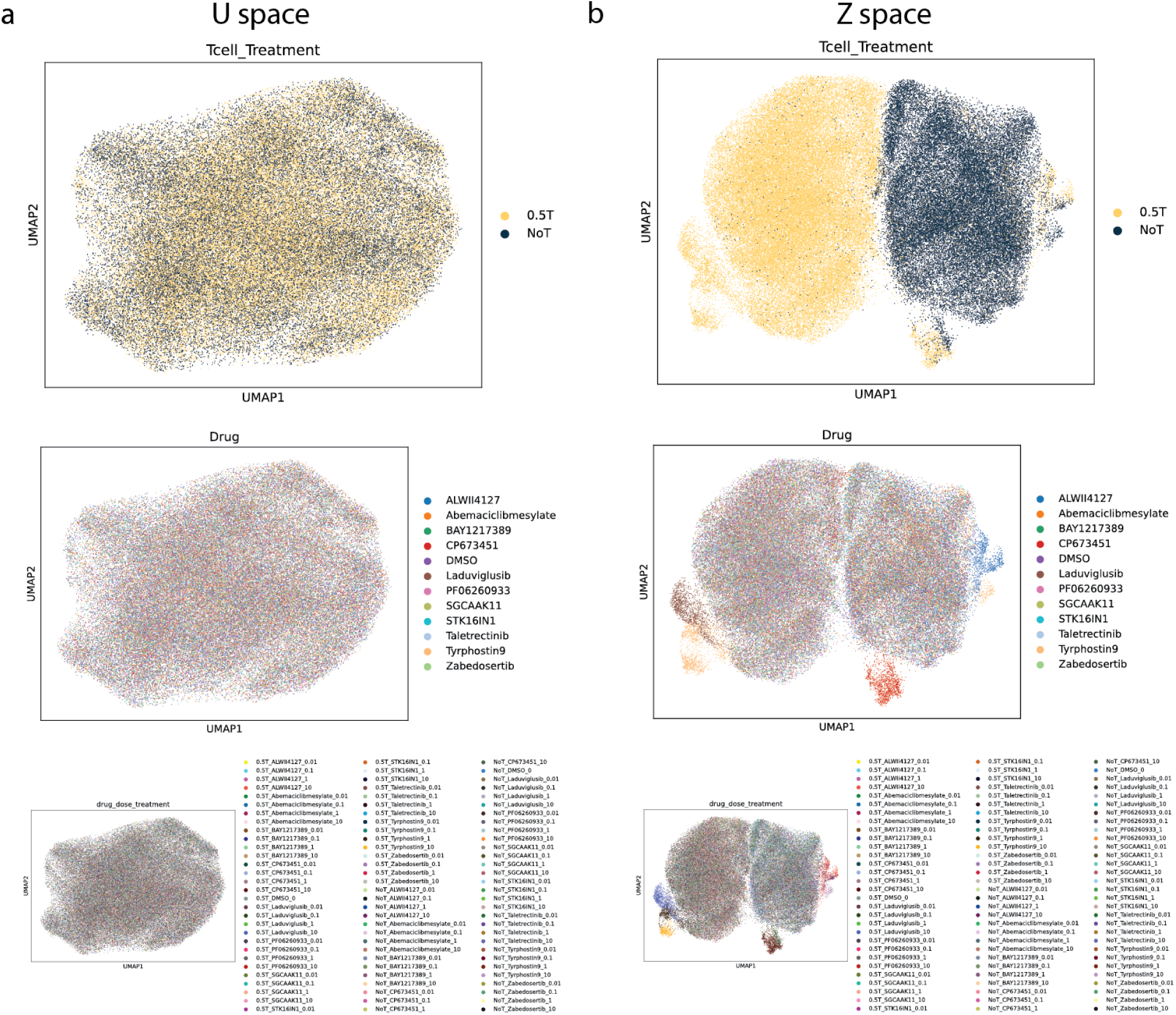
MrVI analysis of PDN single-cell chemical genomics screen for compounds that modulate the T cell-induced program. UMAP visualization of effector-to-target (E:T) ratios in both u space (a) and z space (b), colored by E:T ratios (top), drug treatment (middle), and T cell treatment along with both drug conditions and drug concentration (bottom).

**Supplementary Figure 13.**
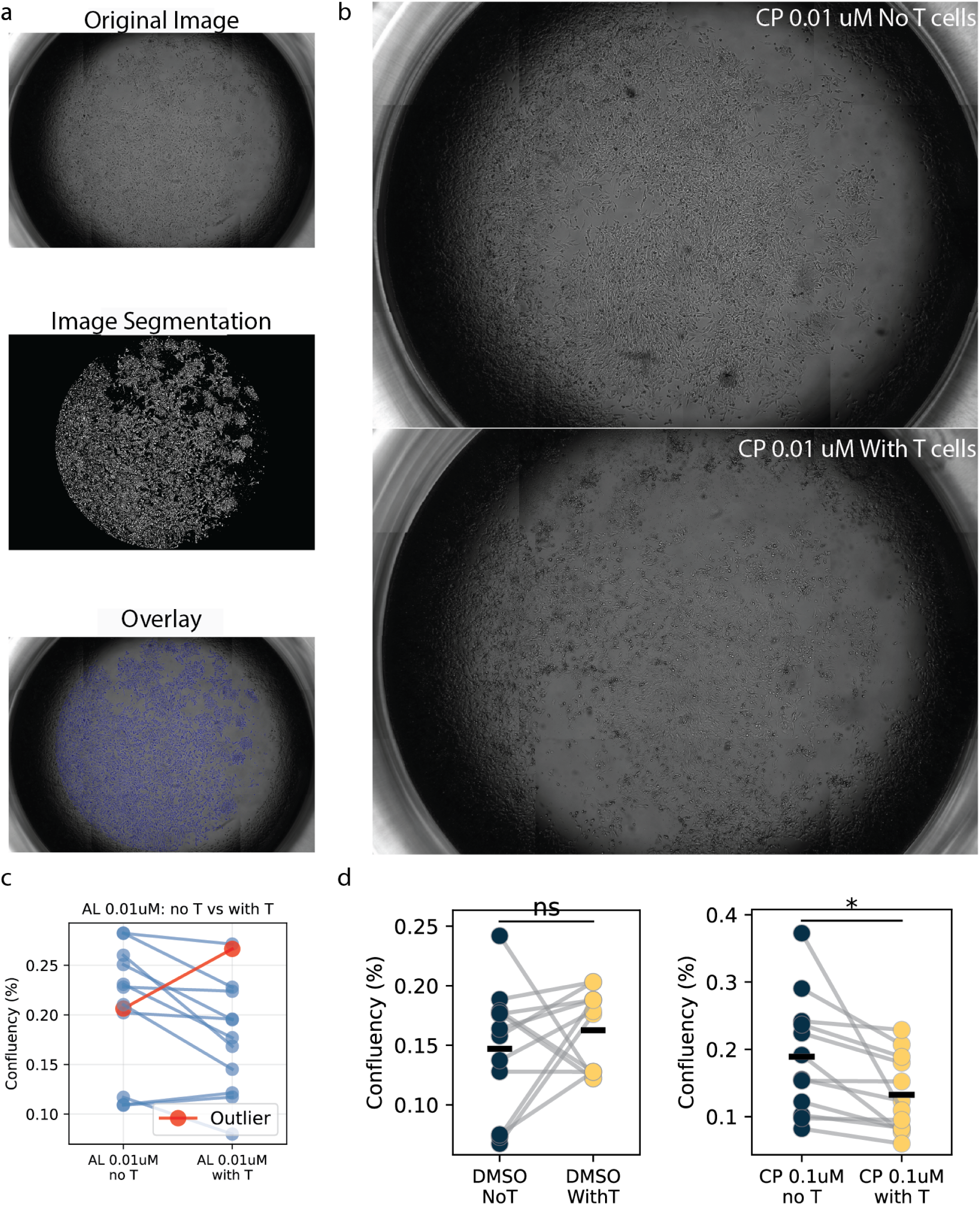
T cell cytotoxic assay analysis. (a) Image segmentation pipeline and overlay applied to brightfield microscopy images taken after 18 hour GBM:T cell co-culture periods for cell confluency analysis. Lower doses of PDGFRA and EphA2 inhibitors improve T cell-mediated killing: b) CP-673451 0.01uM condition shows significant T cell-mediated killing measured by reduced GBM cell confluency. c) Outlier analysis was performed to remove one outlier in the AL 0.01 uM condition. d) Line plots of DMSO controls and CP-673451 0.1uM conditions display significantly reduced mean GBM confluency compared to DMSO control group across all experiments. * indicates statistical significance evidenced by p value < 0.05 for paired t-test results. “ns” indicates results which are not significant. Cell confluency is expressed as a fraction of the well area covered by cells (e.g. 0.20 = 20% of the well is occupied by GBM cells).

**Supplementary Figure 14.**
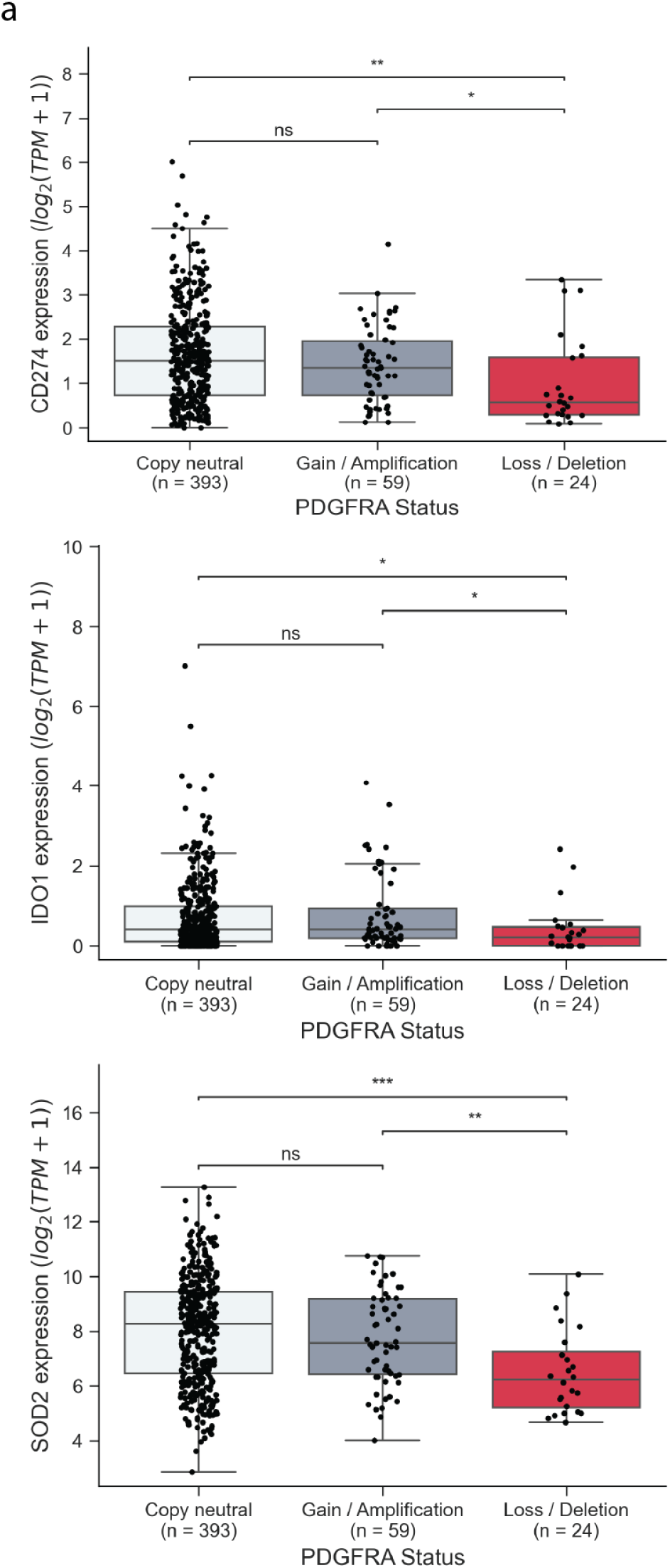
Correlation between PDGFRA copy number and expression of immune evasion genes across bulk RNA samples from the GLASS consortium. Expression of *CD274*, *IDO1*, *SOD2* in bulk RNA-seq data stratified by PDGFRA copy number status across all glioma patients.

**Supplementary Figure 15.**
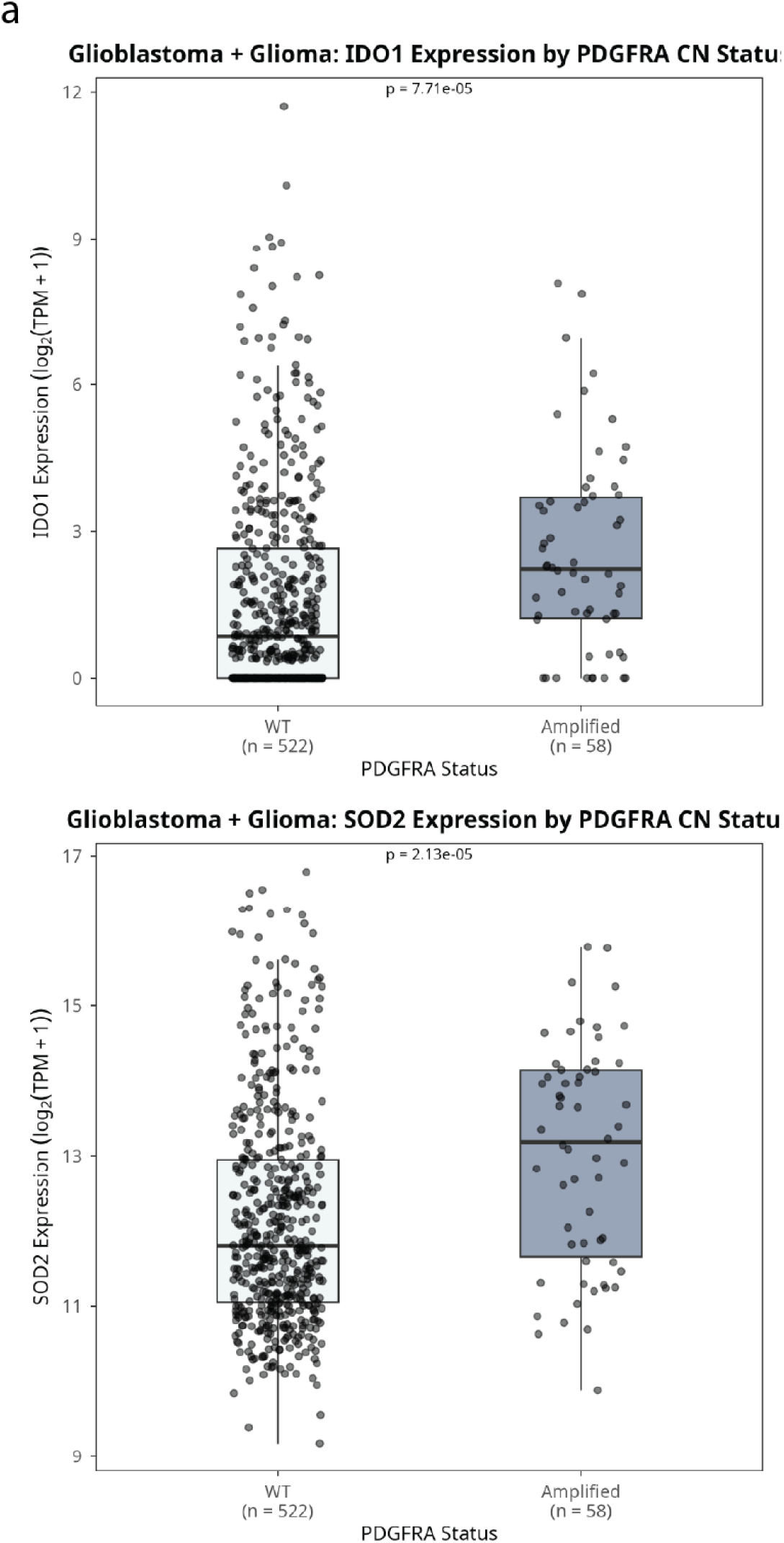
Correlation between PDGFRA copy number and expression of immune evasion genes across bulk RNA samples (Glioblastoma and Glioma) from the TCGA pan cancer atlas^108^. Expression of *IDO1*, *SOD2* in bulk RNA-seq data stratified by PDGFRA copy number status across all glioma patients.

## Methods

### Experimental approaches

#### Cell Culture

The glioblastoma cell lines U87MG were obtained from ATCC and cultured in DMEM (ThermoScientific) supplemented with 10% fetal bovine serum (FBS) and 1% penicillin-streptomycin (P/S), following ATCC-recommended protocols. U87 were maintained in culture and passaged at ∼80-90% confluency using TrypLE (Gibco) according to standard protocols.

Glioma neurosphere cell lines BT112, BT228, and BT333 were provided by the Dana-Farber Cancer Institute (DFCI) Center for Patient Derived Models (CPDM) under a material transfer agreement and maintained as previously described (Touat et al., Nature, 2020). These cells were cultured in NeuroCult™ NS-A Proliferation Medium (StemCell Technologies), supplemented with 0.0002% heparin, 20 ng/mL EGF, and 10 ng/mL FGF (Miltenyi Biotec), under standard conditions of 37°C in a humidified 5% CO₂ incubator. Cells were grown on low-attachment plates coated with Poly(2-hydroxyethyl methacrylate) and passaged using Accutase™ (StemCell Technologies). Poly(2-hydroxyethyl methacrylate) was dissolved at 20 mg/mL in 95% ethanol and incubated at 58°C with shaking at 200 rpm for 2 hours. The solution was then used to coat the plate, which was allowed to air-dry in a biosafety hood under UV light for 2 hours with the fan on. The coated plate can be stored and used for up to one month.

#### Neurosphere Dissociation

Patient-derived GBM neurospheres (cell lines BT112, BT228, and BT333) were dissociated for passaging using Accutase™ (StemCell Technologies, Cat #07920). Cells in culture were gathered and centrifuged at 200 x *g* for 5 minutes. They were then aspirated and resuspended in 1 mL Accutase. They were then placed in a 5% CO_2_ incubator for 7-9 minutes to dissociate neurospheres. After incubation, 5 mL NSA media was added to neutralize the Accutase. The solution was centrifuged at 500 x *g* for 5 minutes, and the pellet was resuspended in an appropriate volume of fresh NSA media. ∼1 million cells were plated in a 10 cm dish in 10 mL NSA media, or ∼200,000 cells/well of a 6-well plate in 3 mL of NSA media/well.

#### Expression of CRISPRi/a systems

The CRISPRi/a system was obtained from McFaline-Figueroa et al^43^. To generate CRISPRi-mediated knockdown cells, lentiviral particles encoding **dCas9-BFP-KRAB** were produced by transfecting HEK293T cells with the plasmid **pHR-SFFV-dCas9-BFP-KRAB** (Addgene, Plasmid #46911) along with the ViraPower Lentiviral Packaging Mix (ThermoScientific). Transfections were performed using Lipofectamine 2000 (ThermoScientific) in Opti-MEM (ThermoScientific), following the manufacturer’s forward transfection protocol.

72 hours post-transfection, the media containing lentiviral particles was collected and filtered through a 0.22 μm Steriflip filtration unit. U87MG glioblastoma cell lines were transduced with the filtered lentiviral supernatant for 48 hours using varying volumes of viral input. Cells were expanded and sorted by fluorescence-activated cell sorting (FACS) to select populations with the highest BFP fluorescence. Transductions were initiated at a multiplicity of infection (MOI) of ∼0.3. To obtain pure populations with consistent dCas9-KRAB expression, cells were expanded and sorted a total of four times.

For the generation of CRISPRa-mediated overexpression cells, the two-component **dCas9-SunTag** system was used. Lentiviral supernatants encoding **dCas9-GCN4-BFP** (pHRdSV40-dCas9-10xGCN4_v4-P2A-BFP, Addgene, Plasmid #60903) and **scFv-GCN4-GFP-VP64** (pHRdSV40-scFv-GCN4-sfGFP-VP64-GB1-NLS, Addgene, Plasmid #60904) were generated as described above. Glioblastoma cells were co-transduced with dCas9-GCN4-BFP at an MOI of ∼0.3 and with scFv-GCN4-GFP-VP64 at an MOI of ∼1. Cells were expanded and sorted by FACS a total of four times based on both BFP and GFP fluorescence to ensure uniform and robust expression across the population.

#### Generation of CROP-seq-OPTI gRNA Libraries

CROP-seq-OPTI gRNA libraries were generated by McFaline-FIgueroa et al.[Citation error]. Protospacer sequences targeting all genes perturbed in this study were sourced from the genome-wide human CRISPRi and CRISPRa v2 libraries developed by Horlbeck et al. (CROP-seq-OPTI, Addgene, Plasmid #106280, engineered to include a CRISPRi-optimized gRNA backbone) were synthesized by Integrated DNA Technologies (IDT). For the kinome-scale screen, oligos were synthesized as a pooled array (CustomArray Inc., Bothell, WA), targeting 522 kinases with a total of 3,165 sgRNAs. Each transcription start site was targeted by five sgRNAs, including both non-targeting and random control sgRNAs.

#### Homology arms for Gibson assembly

- 5’ homology: 5’-ATCTTGTGGAAAGGACGAAACACC-3’
- 3’ homology: 5’-GTTTAAGAGCTATGCTGGAAACAGCATAGCAAGT-3’

Pooled oligonucleotides were amplified via PCR using NEBNext 2X Hi-Fi PCR Master Mix (NEB) and the following primers:

- Forward: 5’-ATCTTGTGGAAAGGACGAAACACCG-3’
- Reverse: 5’-GCTATGCTGTTTCCAGCATAGCTCTTAAAC-3’

Amplification was monitored in real-time using a MiniOpticon system (Bio-Rad) with SYBR Green (Invitrogen), and reactions were terminated before saturation. Amplified products were purified using the NucleoSpin PCR Clean-Up and Gel Extraction Kit (Takara Bio).

The CROP-Seq backbone was transformed with NEB Stable Competent *E. coli* cells, then one E.coli colony was picked and cultured in 100 mL LB broth in the presence of Ampicillin. The CROP-seq-OPTI vector was linearized by sequential digestion with BsmBI and alkaline phosphatase (NEB), with PCR clean-up steps between and after digestion.

Assembly of the vector and sgRNA inserts was carried out using the NEBuilder HiFi DNA Assembly Kit (NEB), with inserts provided at a 20-fold molar excess (55°C for 60 mins). The assembled product was transformed into NEB Stable Competent *E. coli* cells (NEB) via multiple rounds of transformation to ensure sufficient coverage of each sgRNA. Colonies were pooled and cultured in 50 mL Luria broth containing ampicillin at 30°C for 24 hours. Plasmid libraries were extracted using a Qiagen Midiprep Kit. Plasmids were send to both Genewiz and Plasmidsaurus for quality control.

Lentiviral packaging of the gRNA libraries was performed in HEK293T cells transfected with the plasmid library using Lipofectamine 3000 (ThermoScientific) in OptiMEM (ThermoScientific), following the manufacturer’s protocol scaled to 15 cm dishes. Both 48 and 72 hours post-transfection, viral supernatant was collected and filtered through a 0.22 μm Steriflip filter.

To determine viral titers, U87MG glioblastoma cell lines were transduced in 6-well plates with serial dilutions of the filtered supernatant for 72 hours. Cells were then split 1:4 into media with or without 1 μg/mL puromycin. After 96 hours of culture, the multiplicity of infection (MOI) was estimated by comparing cell counts in the presence and absence of puromycin selection.

For large-scale screens, 1.2 million cells were seeded in a 6-well plate and transduced at an MOI of ∼0.3 with spin infection (800g for 60 mins in RT). 72 hours post-transduction, cells were pooled and transferred to two 15 cm dishes with 1 μg/mL puromycin and maintained under selection. Cells were seeded for T cell treatment experiments between 10 to 14 days post-transduction.

#### Generation of NY-ESO-1+ GBM Cells

U87MG glioblastoma/PDN cells were engineered to stably express the tumor antigen NY-ESO-1 (CTAG1B). A plasmid encoding NY-ESO-1 was obtained from OriGene Technologies (RC213318). The NY-ESO-1 insert was PCR-amplified and cloned into a lentiviral backbone containing a blasticidin resistance cassette (Addgene plasmid #52962) using Gibson assembly. Primers were designed to append vector-homology overhangs to the NY-ESO-1 amplicon, and assembly was performed using a 2-fold molar excess of insert at 55 °C for 60 min. Lentiviral particles were produced by transfecting HEK293T cells with the resulting plasmid using the Lipofectamine 3000 transfection kit (Thermo Fisher Scientific). Cells were transduced with the viral supernatant and selected with 10 μg/mL blasticidin for 3 days. Cells were subsequently maintained in culture with 1 μg/mL blasticidin to ensure stable integration and continued NY-ESO-1 expression. Overexpression efficiency is validation via flow cytometry with NY-ESO-1 antibody (Santa Cruz Cat #E978).

Primers (PCR amplification and vector overhangs)

- NYESO1_Forward: GCGATCGCCATGCAGGCCGAAG
- NYESO1_Reverse: GCCCTGAGGGAGGCTGAGCCAAAAAC
- Lenti_backbone_Forward: GGATCCGGCGCAACAAACTTCTCTCTGC
- Lenti_backbone_Reverse: GGTGGCAGCGCTCTAGAACCGG
- NYESO1_Forward with overhang: CCGGTTCTAGAGCGCTGCCACCGCGATCGCCATGCAGGCCGAAG
- NYESO1_Reverse with overhang: GCAGAGAGAAGTTTGTTGCGCCGGATCCGCCCTGAGGGAGGCTGAGCCAAAAAC

#### Generation of HLA-A*02:01+ GBM cells

PDN cells were engineered to stably express HLA-A*02:01 to immune matched with the T cells. A plasmid encoding HLA-A*02:01 was obtained from (Addgene, Plasmid #108213). The HLA sequence was PCR-amplified using the primers listed below. The resulting amplicon was cloned into a lentiviral backbone containing a puromycin resistance cassette (Addgene plasmid #108100) using Gibson assembly. Primers were designed to append vector-homology overhangs to the HLA fragment, and assembly was performed using a 2-fold molar excess of insert at 55 °C for 60 min. Lentiviral particles were produced by transfecting HEK293T cells with the resulting plasmid using the Lipofectamine 3000 transfection kit (Thermo Fisher Scientific). PDN cells were transduced with the viral supernatant and selected with 1 μg/mL puromycin for 4 days. Overexpression efficiency is validation via flow cytometry with HLA-A2 antibody (Biolegend, 343304).

- HLAA0201_Forward: ATGGCGCCCCGAACCCTCGTC
- HLAA0201_Reverse: CACTTTACAAGCTGTGAGAGACACATCAGAGCCC
- Lenti_backbone_Forward: GGCAGCGGCGCCACCAACT
- Lenti_backbone_Reverse: GGTGGCGGATCCCGCGTCAC
- HLAA0201_Forward with overhang: GTGACGCGGGATCCGCCACCATGGCGCCCCGAACCCTCGTC
- HLAA0201_Reverse with overhang: AGTTGGTGGCGCCGCTGCCCACTTTACAAGCTGTGAGAGACACATCAGAGCCC

#### Verification of Protein Overexpression

Flow cytometry was used to confirm the expression of HLA-A0201 and NY-ESO-1in the engineered GBM cell lines. Cells were stained with fluorescent antibodies specific to HLA-A0201 and NY-ESO-1, along with appropriate controls. Data was analyzed using FlowJo software to compare protein expression between engineered and non-engineered cells.

#### Antibody staining of NY-ESO-1 antigen

Cells were first dissociated or trypsinized using the procedure mentioned above. In single-cell form, cells are washed twice with FACS buffer (2%FBS in PBS) and centrifuged at 500 x *g* for 5 minutes at 4°C in between each wash. Cells were then suspended in 100 µL of 4% PFA for fixing and incubated at 4°C for 15 minutes. After two washes, cells were resuspended in 100 µL of 0.1% TritonX for permeabilization and incubated for 15 minutes at room temperature. After two more washed, cells were stained with NY-ESO-1 antibody (Santa Cruz, Cat #E978) in a ratio of 1:25 for 30 minutes at 4°C.

#### Antibody staining of HLA-0201 surface protein

Cells are first to be dissociated using the procedure mentioned above, and then washed twice. Cells are then stained live with with the HLA-A2 antibody (BioLegend, Cat #343303) in a ratio of 1:25 for 30 minutes at 4°C.

#### T cell-mediated killing assay

For the cytotoxic T cell killing assay, terminally differentiated human CD8⁺ T cells specific to NY-ESO-1 were obtained from Charles River Laboratories (ASTC-1093). Upon receipt, T cells were thawed following the supplier’s protocol and used immediately.

300k cells of U87MG NY-ESO-1+ cells were seeded in RPMI media with 10% FBS in each well of a 6 well plate with 1 ng/ml IFN-γ overnight. U87MG-T cell co-culture at various ratios was performed in complete T cell media composed of RPMI 1640, supplemented with 10 mM HEPES, 10 mM L-glutamine, 10% FBS, 0.34% β-mercaptoethanol, and 1% penicillin-streptomycin^138,139^. After 18 hours of co-culture, media with T cells were removed, and U87MG NY-ESO-1+ cells were detached using TrypLE (Gibco). T cells and GBM cells were next subjected to 10X single cell genomics and Sci-Plex protocol^140^, respectively.

Patient-derived GBM neurospheres (PDN) were dissociated as described above, and 50,000 cells were seeded into ultra-low attachment 96-well plates. After a 24-hour recovery period, small molecule kinase inhibitors were added at final concentrations of 10, 1, 0.1, or 0.01 µM, prepared from 10 mM DMSO stock solutions and diluted in complete NS-A medium. DMSO was used as a vehicle control at a final concentration of 0.1% (v/v). Cells were incubated with inhibitors for 24 hours at 37°C and 5% CO₂, followed by the addition of IFN-γ to a final concentration of 1 ng/mL for an additional 24 hours. T cells were added at a ratio of 0.5 T cells per PDN for 18 hours of co-culture. Cells were then harvested for the Sci-Plex protocol^140^.

#### Nuclei Hashing and Fixation

Nuclei hashing and fixation procedures were adapted from protocols by Srivatsan et al^140^ and Sziraki et al^141^. Adherent cells were trypsinized with TrypLE (Gibco) while GBM neurospheres were dissociated. After washing with ice-cold 1x PBS, cells were lysed using EZ Lysis Buffer (Sigma) supplemented with 1% diethyl pyrocarbonate (Sigma), 0.1% Superase•In RNase Inhibitor (Thermo Fisher), and 500 fmol of hashing oligo. Following lysis, nuclei were fixed by adding 1.25% formaldehyde in 1.25x PBS, resulting in final concentrations of 1% formaldehyde and 1x PBS, and incubated on ice for 10 minutes. Nuclei were then pooled into a plastic reservoir and transferred to 50 mL conical tubes for centrifugation at 650 x *g* for 5 minutes at 4°C. The supernatant was removed, and nuclei were washed once with nuclei suspension buffer (NSB), composed of 10 mM Tris-HCl (pH 7.4), 10 mM NaCl, 3 mM MgCl₂, 1% Superase•In, and 1% Ultrapure BSA (0.2 mg/mL, New England Biolabs). Nuclei were then resuspended in NSB, slow-frozen in 10% DMSO, and stored at −80°C until further processing by sci-RNA-seq.

#### Library Preparation and Sequencing

Frozen, hashed nuclei were thawed and processed using a three-level combinatorial indexing protocol adapted from Cao et al.^142^, Martin et al.^143^, and Sziraki et al^141^. Nuclei were spun down, resuspended in NSB, and briefly sonicated for 12 seconds at low power using a Bioruptor. After counting, 21 μL of nuclei were dispensed into each well of 96-well low-adhesion PCR plates. Each well received 2 μL of 10 mM dNTPs, 2 μL of 100 μM indexed oligo-dT primers, 2 μL of 100 μM indexed random hexamer primers, and 14 μL of a reverse transcription master mix containing 100 mM DTT, RNaseOUT, 5× SuperScript IV buffer, and SuperScript IV reverse transcriptase. Reverse transcription was performed using a gradual temperature ramp. After reverse transcription, all nuclei were pooled and redistributed into new 96-well plates (10 μL per well) for ligation. Each well was supplemented with 8 μL of indexed ligation primers, 4.8 μL of a 3:2 mix of T4 ligase buffer and T4 ligase (NEB), and 9.4 μL of nuclei buffer with BSA (NBB), which had the same composition as NSB but with BSA.

Ligation reactions were incubated at 25°C for 1 hour. Nuclei were then pooled, washed with NBB, and redistributed at a concentration of 1,500 nuclei in 5 μL of NBB per well. Cells can be stored for future use. For second strand synthesis, 5 μL of synthesis mix—comprising 60% Qiagen Elution Buffer, 27% NEB Second Strand Synthesis Buffer, and 13% NEB Second Strand Enzyme Mix—was added to each well, and the reaction was carried out at 16°C for 3 hours. Tagmentation was then performed by adding 1/50 μL of N7-adaptor-loaded Tn5, followed by incubation at 55°C for 5 minutes and quenching with DNA binding buffer (Zymo) at room temperature for 5 minutes. Double-stranded DNA was purified with a 1X SPRI bead cleanup step carried out directly within the 96-well plates. The resulting DNA was subjected to USER enzyme digestion using a solution of 80% nuclease-free water, 10% 10X rCutsmart buffer, and 10% USER enzyme (NEB). DNA was eluted into a clean 96-well plate using Buffer EB.

To each well, 16 μL of eluted product was mixed with 2 μL each of P5 and P7 PCR primers in a well-specific indexed combination, along with 20 μL of 2X NEBNext PCR Master Mix (NEB). PCR was used to amplify libraries and incorporate sequencing adapters. Final PCR products were pooled, and SPRI bead purification was performed with a 0.7X cleanup for cDNA libraries and a 1X cleanup for hash libraries.

#### 10x Genomics Library Preparation

T cells were processed for single-cell RNA sequencing using the 10X Genomics platform according to the manufacturer’s protocols. 4000 cells were loaded on a Chromium X using the Chromium Next GEM Single Cell 3’ Kit v3.1 (10x Genomics, PN-1000123). After reverse transcription and cleanup, cDNA libraries were generated as per the 10x user guide. Construction of final gene expression (GEX) libraries was performed using the Library Construction Kit (10x Genomics, PN-1000190) and Dual Index Kit TT set A (10x Genomics, PN-1000215) according to the user guide. The fragment size distribution of cDNA and final sequencing-ready GEX libraries was assessed using a TapeStation 2200 system (Agilent) with TapeStation D5000 reagents.

#### scRNA library sequencing

Library concentrations were measured using a Qubit fluorometer (Invitrogen), and quality was assessed via TapeStation DNA D1000 assays. Sequencing was performed on an Illumina NextSeq 550 platform and Element Biosciences AVITI system, following the manufacturer’s protocols. A 2% PhiX spike-in was included to monitor sequencing quality.

#### TCR CDR3 sequences enrichment from 10x Genomics 3’ library

##### Nested PCR enrichment approach

To capture single-cell TCR clonotype information, we adapted an established protocol^144^. The process involves three qPCR steps: (1) the first step begins with 43 pooled TCRB primers (pooled at 100µM, diluted to 10µM) and a truncated read 1 primer (10µM) (2 µl cDNA, 1 µl of each forward and reverse primer and 12.5 µl 2X NEBNext Master Mix, 0.5 µl SYBR and 8 µl water). (2) The second step follows the same structure, but substitutes the forward primer with 43 pooled TCRB primers containing R2 sequences (10µM), and 1 µl of purified PCR product from step one as the template. (3) The third step involves indexed 10µM TruSeq P5 primers and indexed Nextera P7 primers, with 2 µl of purified PCR product from step 2 scaled to a 50µL reaction. All PCR steps were stopped before the plateau phase, and the PCR products were cleaned with 0.8x SPRI beads and eluted in 50 µl.

The PCR cycling conditions are as follows: initial denaturation, 98 °C for 3 min; denaturation, 98 °C for 15 s; annealing, 62 °C (72 °C for qPCR step 3) for 20 s; extension, 72 °C for 1 min (with an extra 15 s post plate read); repeat of the denaturation step to the extension step before the plateau phase; final extension, 72 °C for 3min. We further provide the full spatial TCR primer sequences in Supplementary Table 15.

##### RhPCR enrichment approach

Multiplex rhTCR amplification was adapted from Liu et al. (2022)^145^ and Li et al. (2019)^146^. The protocol consists of two qPCR steps. In the first step, TRA and TRB loci were amplified using 69 rhPCR-modified primers together with modified P5 and P7 primers and RNase H2. Reactions (24 µL) contained 2 µL cDNA, TRXV primer pool (0.05 µM per primer), RNase H2 (0.5 mU/µL), modified P5 and P7 primers (2 µM each), 2x SYBR® Green BioRad Master Mix (1708880) and water. qPCR was stopped prior to the plateau phase, followed by a final extension, and products were purified using 0.8x SPRI beads and eluted in 20 µL.

In the second step, full-length adapters were completed using unmodified P5 and P7 primers and NEBNext High-Fidelity 2x PCR Master Mix with 5 µL of PCR product from step 1 in a 50 µL reaction. qPCR cycling was terminated before plateau. Libraries were purified with 0.8x SPRI beads, assessed by Bioanalyzer, and prepared for sequencing. We further provide the full TCR primer sequences in Supplementary Table 16.

The PCR cycling conditions are as follows for qPCR step 1: initial denaturation, 95 °C for 5min; denaturation, 96 °C for 20 s; annealing, 60 °C for 4 min; extension, 72 °C for 1 min (with an extra 15 s post plate read); repeat of the denaturation step to the extension step before the plateau phase; final extension, 72 °C for 3 min. For qPCR step 2 the conditions are: initial denaturation, 98°C for 30 s; denaturation, 98 °C for 10 s; annealing, 72 °C for 20 s; extension, 72 °C for 1 min (with an extra 15 s post plate read); repeat of the denaturation step to the extension step before the plateau phase; final extension, 72 °C for 3 min.

#### Cyctotoxic killing assay

For a full 96-well plate at a final concentration of 2.5 µg/mL, 160 µL of thawed Recombinant Human Laminin-521 (Thermofisher, Cat #A29248) stock solution was added to 6.24 mL of DPBS containing calcium and magnesium ions (Thermofisher, Cat #14040133). The solution was thoroughly pipette-mixed and 64 µL was added to each well. The plate containing solution was incubated at 37°C for 2 hours, then removed and parafilmed for storage at 4°C until use within 2-3 days. The laminin solution was aspirated immediately before seeding of dissociated GBM PDN cells.

24 hours after seeding 25,000 patient-derived GBM cells (BT333) per well on laminin-coated plates, media was replaced with a serial dilution of drugs CP-673451 and ALW-II-41-27 at concentrations 0.1uM, and 0.01uM vertically. This was repeated for the bottom half of the plate that will contain T cells. Drugs were dissolved at a final concentration of 0.1% DMSO. 24 hours after drugging GBM cells, IFN-γ was added to each well in a final concentration of 1 ng/mL. After 24-hour IFN-γ pretreatment, 100 µL of 25,000 Anti NY-ESO-1 CD8+ T cells (Charles River, ASTC-1093) suspended in complete RPMI media were added to the bottom half of the 96-well plate.

After 18 hours of GBM-T cell co-culture, Brightfield images were captured using a Zeiss Axio Observer microscope equipped with an AxioCam 205 color camera to control image acquisition in ZEN Pro software. Illumination was provided by an X-Cite fluorescence illumination system (Excelitas Technologies). Images were captured at 5X magnification. For each well, an identical 4x4 tiled grid was applied in ZEN Pro to capture the majority of the well area. Z-axis focal positions were individually optimized for each well to capture the viable GBM cell populations.

#### Computational analysis

##### Processing of raw sequencing results and generation of count data matrix

Sequences were demultiplexed using bcl2fastq (Illumina) filtering for reads with RT and ligation barcodes within an edit distance of 2 bp. PolyA tails were trimmed using trim-galore (https://github.com/FelixKrueger/TrimGalore) and reads were mapped to the human hg-38 transcriptome using STAR^147^. After alignment, reads were filtered by alignment quality and duplicates were removed if they mapped to the same gene, the same barcode and the same unique molecular identifier (UMI) or if they met the first 2 criteria and the UMI was within an edit distance of 1 bp. Reads were assigned to genes using bedtools^148^. 3’ UTRs were extended by 100 bp in the gene model to account for short 3’ UTR annotations to minimize genic reads labeled as intergenic. A knee plot was used to set a threshold above which a combinatorial cell barcode confidently corresponded to a cell. UMI counts for cell barcodes that pass this threshold were aggregated into a sparse matrix format, followed by the creation of a cell dataset object using Monocle3^149–151^.

##### Hash and sgRNA assignment

Sci-Plex hashes and sgRNA transcripts derived from CROP-seq were isolated from demultiplexed reads. Hashes were assigned as previously described^140^. Briefly, reads were considered hashes if (1) the first 10 bp of read 2 matched a hash in a hash whitelist within a Hamming distance of 2 and (2) contained a poly-A stretch spanning the 12–16 bp region of read 2. For sgRNA assignment, reads were considered CROP-seq derived if the bases spanning position 24–42 matched a sgRNA in a sgRNA whitelist within a hamming distance of 2 and (2) a TGTGG sequence at position 3–7 of read 2. Duplicated reads were collapsed by their UMIs arriving at hash and sgRNA UMI counts for each nucleus in our experiment. Finally, we tested whether a particular nucleus was enriched for one or more hash or sgRNA as described in^140^ for sci-Plex hashes.

##### Data preprocessing and dimensionality reduction

To filter out cells with low quality, knee plots of UMI counts per cell were drawn to determine a reasonable threshold. Cells with fewer than 500 UMIs were removed. Hash and sgRNA were assigned to cells based on their corresponding cell barcodes, disregarding P7 sequences. Doublets were detected using *Scrublet^152^*.

Then, the data were processed with an initial round of dimensionality reduction and *Leiden^153^* clustering to remove T cell contamination. Specifically, top 5000 highly dispersed genes were selected with *estimateDispersionsForCellDataSet* in Monocle3^149–151^ package and used for principal component analysis (PCA)(15 PCs). To generate a 2D visualization of the data with UMAP, we specified the parameters in *reduce_dimension* function as follows: *umap.n_neighbors=20L*, *umap.min_dist=0.1*, *max_components=2*, *umap.fast_sgd=FALSE*. With a resolution of 2e-4, cells were partitioned into clusters, and cells with obvious expression of canonical T cell lineage markers (*CD8A and CD3D*) were filtered out.

For the CRISPRi dataset, a total of 182,652 cells were retained, with a mean UMI count of 4,577 and a median UMI count of 2,865. The median number of cells per perturbation was 85.5, 111, 75, and 59 for E:T ratios of 0, 0.25, 0.5, and 1, respectively.

For the CRISPRa dataset, a total of 317,405 cells were retained, with a mean UMI count of 5,764 and a median UMI count of 4,503. The median number of cells per perturbation was 156, 118, 143, and 97 for E:T ratios of 0, 0.25, 0.5, and 1, respectively.

#### Differential gene expression analysis

All differential gene expression analyses were performed using *fit_models* function in the Monocle3^149–151^ package, which fits generalized linear models (GLM). By default, it uses quasi-poisson distribution to model the gene expression value, represented by raw read counts normalized by size factor.

To investigate the effect of T cell treatment on gene expression profiles of GBM cells, we defined a signature of T cell dose response genes on non-targeting control (NTC) cells from CRISPRa and CRISPRi dataset respectively. For every gene that was expressed in at least 5% of cells in the NTC dataset, we fit a GLM on two coefficients, T cell treatment conditions and biological replicates, with default settings of *fit_models* function. P values generated from the models were corrected collectively for multiple hypothesis testing using the Benjamini-Hochberg method. As a result, a list of T cell dose response genes were selected with adjusted p value < 0.05 and abs(normalized effect size) >0.25.

Additional differential expression analyses were performed using similar frameworks but with different cell subsets and model formulations. These analyses identified (1) genotype-associated genes in untreated cells, (2) genotype-dependent T cell dose–response genes within each treatment condition, and (3) treatment dose–associated genes within each genotype.

#### Calculation of aggregated gene scores for proliferation, resistance and APM gene programs

Antigen-presentation machinery scores were calculated as in Srivatsan et al^140^. Specifically, for each cell, raw expression counts for genes within the program were extracted from the count matrix and normalized using size factors. The normalized counts were then aggregated by gene program and log-transformed with a pseudocount of 1. The scores were then used for differential expression test as described previously to determine which kinases significantly modulate antigen-presentation machinery.

#### MrVI model training

To systematically evaluate and compare the effects of perturbation across different T cell treatment conditions on transcriptomic profiles, we employed MrVI^88^, a deep generative framework designed for integrating and comparing multi-sample and multi-batch single-cell transcriptomic data. Briefly, MrVI learns two latent spaces from a given dataset, where *U*-space is a sample-unaware space designed to capture high-level cell state heterogeneity, while *Z*-space further incorporates sample-of-origin information. This hierarchical architecture enables us to perform robust sample stratification at a single-cell resolution.

Here, we trained a MrVI model on CRISPRa and CRISPRi dataset separately with default settings. Prior to training, we applied stringent subsetting to ensure high-confidence model performance and interpretability. First, a list of high-impact kinases was defined based on the number of T cell dose response genes they modulate in untreated cells. Kinases of interest were selected based on knee plot threshold. The dataset was then subset in three steps: 1) Cells were required to have a total sgRNA UMIs per cell > 3 and a ratio of the most abundant sgRNA UMIs to total CROP-seq UMIs >0.3. 2) The dominant sgRNA should be targeting a kinase in the curated list with more than 25 cells, and non-targeting control (NTC) cells were down-sampled to match the abundance of the second most frequent sgRNA. 3) Cells were required to have a total hash UMIs per cell > 2 and the ratio of the most abundant hash UMI to the next abundant hash UMI > 2. In addition, genes were restricted to T cell induced cancer cell responses genes only (Supplementary Tables 1, 2).

During model training, the unique E:T ratio -genotype combination was specified as the sample key, while the batch key corresponded to biological replicates. The model was trained at a maximum of 400 epochs, which was sufficient given the validation evidence lower bound (ELBO). We visualized the *U*-space and *Z*-space by UMAP embeddings as implemented in Scanpy^154^ (*scanpy.pp.neighbors* and *scanpy.tl.umap*) with default parameters. To evaluate the similarity of perturbation effects, MrVI defines a distance metric which measures Euclidean distances between counterfactual cell states in *Z*-space for a given cell in *U*-space. In this context, for any cell in its original dosage-genotype condition, a counterfactual cell state represents the predicted transcriptomic profiles that the same cell would exhibit under an alternative condition. Formally, the latent representation is defined as

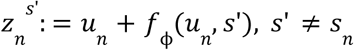

where *f*_ϕ_ is the learned effect of the condition. This approach enables us to calculate a pairwise distance matrix between all dosage-genotype combinations for any given cell. Aggregating across all untreated cells yields a unified distance matrix that summarizes perturbation effect at a single-cell resolution.

To quantify how kinase perturbations altered the expected E:T ratio ordering in MrVI latent space, we first computed pairwise local sample distances from the MrVI model and converted them to a distance matrix between all kinase–E:T ratio samples. For each sample, we calculated the mean distance to every E:T ratio group (0, 0.25, 0.5, 1) and identified cases where the sample was closer, on average, to a different E:T ratio group than its own. For these "misaligned" samples, we compared the distribution of distances to the own dosage group versus the closest other group using a one-sided Mann–Whitney U test. P-values were Bonferroni-corrected across all tested samples, and only samples with corrected p < 0.05 were retained. The figure plots significant samples as points at their own dosage along the x-axis and kinase on the y-axis, with point size and color encoding Bonferroni-corrected significance and arrows indicating the direction and relative magnitude of the shift toward the closest dosage group.

#### Decipher trajectory analysis

To characterize cell states and their distribution across perturbations and conditions, we applied Decipher[Citation error] to the GBM cells. The Decipher algorithm was used to learn a low-dimensional embedding that captures the differentiation of T cells while preserving biological heterogeneity across different E:T ratios. We filtered NTC cells hashing quality metrics (total_hash_umis_per_cell_ID > 3 & top_to_second_best_ratio > 2), then restricted to genes identified as T cell induced cancer response genes like MrVI from an GLM differential expression analysis (q-value < 0.05, |normalized_effect| > 0.25). We configured Decipher with (*decipher_config = dc.tl.DecipherConfig()*) and applied it to the raw count data. Training was performed using *dc.tl.decipher_train()* with *plot_every_k_epochs=5* to visualize the embedding during training. Both the Decipher embedding was rotated and trajectories were constructed from low (0) to high (1) effector-to-target (E:T) ratios.The resulting two-dimensional embedding provided a biologically interpretable representation of the NTC GBM cell states.

Gene module analysis was performed with SCORPIUS^155^. Gene importance scores along the pseudotime were computed with permutation-based testing (gene_importances, 10 permutations, 6 threads), and p-values were adjusted by Benjamini–Hochberg; genes with q-value < 0.05 were retained.

To align the trajectories for both CRISPRi and CRISPRa, both dataset were concatenated and restricted to shared genes, and filtered on hashing quality (total_hash_umis_per_cell_ID > 3, top_to_second_best_ratio > 2). Decipher latent space was rotated to align the primary axis with E:T ratio and a secondary axis with perturbation type. We defined Decipher trajectories separately for CRISPRi and CRISPRa for Untreated (E:T=0) to 1:1 (E:T=1), which were then used by *dc.tl.trajectories* to compute modality-specific ratio-response paths in the Decipher latent space. Finally, for each perturbation (CRISPRi or CRIPSRa), the distribution of decipher_time for each guide was statistically compared to NTC using Kolmogorov–Smirnov and Mann–Whitney U tests and significant guides exported and visualized via density plots overlaid with Decipher trajectory.

#### Validation of T cell induced cancer cell response signatures in the GLASS cohort

To validate T cell induced cancer cell response genes in brain tumors, we analyzed transcriptomic and clinical data from the GLASS (Glioma Longitudinal AnalySiS) consortium^156^. T cell induced cancer cell response genes with a false discovery rate–adjusted q value < 0.05 were ranked by normalized effect size; the 100 most strongly up-regulated and 100 most strongly down-regulated genes defined the “up-regulated” and “down-regulated” T cell dose–response signatures. Since very limited glioblastoma samples in GLASS are annotated with PDGFRA loss/deletion, we included all glioma samples of various disease subtypes. Patients with multiple samples but have different copy number calls for PDGFRA were filtered out, resulting in a final cohort of 476 samples. Bulk RNA-seq TPM values were extracted from the gene-level expression matrix, *log*_2_ (*TPM* + 1)-transformed and merged with associated clinical annotations using matched case barcodes and sample types. These scores were compared across all GLASS tumors to assess the robustness and clinical relevance of the experimental T cell induced cancer cell responses signatures in an independent, longitudinal glioma cohort.

#### Microscope Image Quantification via Python Analysis

Cell confluency was quantified using a custom Python analysis pipeline implemented with OpenCV (cv2) and NumPy. For each microscope image, grayscale images were loaded using cv2.imread() with the IMREAD_GRAYSCALE parameter. To exclude the dark well edge from analysis, a circular mask was created centered at the image midpoint with a specified pixel radius. The total well area in pixels was calculated as the sum of all pixels within the circular mask. To account for uneven illumination, adaptive Gaussian thresholding (block size 51, C=8) was applied with binary inversion to segment dark cell bodies from the bright background. The resulting mask was refined using morphological opening (3x3 kernel) to remove noise. Cell confluency was calculated as the proportion of well area covered by cells: confluency = cell area pixels / well area pixels.

To compare the cell confluency results between experimental conditions, pairwise comparisons were systematically performed for each drug at each concentration, comparing conditions with T cells versus without T cells across all independent experiments. Statistical comparisons were performed using paired t-tests implemented with scipy.stats.ttest_rel() with alternative=’greater’ to test the directional hypothesis that confluency differed between T cell versus No T cell treatment.

#### Cell-Cell Communication Analysis

Cell-cell communication between GBM and T cells was analyzed using NicheNet^75^ v2.0 in R. Single-cell RNA-seq data were converted from Monocle3 objects to SingleCellExperiment format, and gene symbols were updated to HUGO nomenclature using ‘convert_alias_to_symbols()’ and ‘make.names()’. Datasets were harmonized by identifying common genes and combining with ‘cbind()’. Cells were filtered to two conditions (E:T = 0: GBM alone; E:T = 1: 1:1 T cell:GBM co-culture) and balanced to a 2:1 GBM:T cell ratio via stratified sampling. Genes expressed in < 3% of cells were filtered, and expression data were log-normalized using ‘logNormCounts()’.

Human NicheNet v2 ligand-receptor networks and ligand-target matrices were downloaded from Zenodo (DOI: 10.5281/zenodo.7074291, 10.5281/zenodo.10229222). Expressed genes were defined as those detected in ≥10% of cells per cell type. Differentially expressed genes in receiver cells were identified using Wilcoxon rank-sum tests with Benjamini-Hochberg correction (adjusted p < 0.05, log2FC > 0.25); if <20 genes met criteria, the top 100 upregulated genes were used. Bidirectional analyses (GBM→T cells and T cells→GBM) identified potential ligands expressed in sender cells with cognate receptors in receiver cells. Ligand activities were calculated using ‘predict_ligand_activities()’, ranking ligands by area under the precision-recall curve (AUPR). The top 20 ligands were selected, and ligand-target links were inferred using ‘get_weighted_ligand_target_links()’ (n=200 targets per ligand).

Network visualizations were generated using ggraph and igraph with three layouts: circular, stress-minimization, and Fruchterman-Reingold force-directed. Edge thickness and transparency were scaled to ligand activity scores.

#### TCR enrichment computational methods

Sequencing data were demultiplexed and quality controlled using bases2fastq (Element Biosciences). TCR reads were processed with TRUST4 v1.0.13 to align reads, identify CDR3 sequences and V(D)J usage, and associate clonotypes with cell barcodes. TRUST4-generated barcode AIRR files were used for downstream analyses. To increase confidence of identified clonotypes, barcodes were filtered requiring a minimum consensus (UMI) count of ≥2 per assigned CDR3.

#### Single-Cell RNA-Seq Analysis of T Cells

Single-cell RNA-seq data from T cells cultured alone at four effector:target (E:T) ratios were processed using Scanpy^154^. Quality-control metrics (genes per cell, total UMI counts, mitochondrial and ribosomal percentages) were computed; cells with low gene (>200) or high mitochondrial content were removed (<10), genes captured by less than 3 cells were removed and doublets were identified and excluded using Scrublet^152^. The mean UMI count was 5,723. After quality control, 5,884 cells were retained, including 2,994 untreated cells and 426, 943, and 1,524 cells from the 0.5T:1GBM, 1T:1GBM, and 2T:1GBM conditions, respectively. Counts were normalized per cell, log-transformed, and 5,000 highly variable genes were selected and used for downstream analysis. A k-nearest-neighbor graph was constructed on the highly variable genes, followed by UMAP embedding and Leiden community detection. A non–T-cell cluster was removed based on marker expression and QC metrics, and the graph and UMAP were recomputed on the filtered T-cell population.

#### Spatial Transcriptomic Analysis of Public Mouse GBM Datasets

Spatial transcriptomic data from a publicly available 10x Visium mouse glioma sample (GSM7839621) were analyzed using Scanpy. Spots annotated as tumor were subset, and T cell induced programs were imported from a prior R-based differential expression analysis of non-targeting control CRISPRa T cell cocultures; genes with a q-value <0.05, and top 30 normalized effect were selected, mapped to mouse symbols, and intersected with the Visium gene set. Per-spot T cell induced cancer cell response signature scores and MHC class I antigen-processing/presentation scores were then computed with sc.tl.score_genes using the filtered T cell induced cancer cell response signature and a curated MHC class I gene list, respectively. To stratify spatial regions by T cell infiltration, tumor spots were classified as "T cell infiltrated" or "No T cells" based on detectable expression of TCR chain genes (*Trac*, *Trbc1*, *Trbc2*), and spatial feature plots of Cd3g and Trac were generated. T cell dose–response and MHC class I scores were compared between infiltrated and non-infiltrated tumor regions using violin plots and two-sided Mann–Whitney U tests. For T cell–infiltrated spots, we further quantified the relationship between T cell dose–response and TCR marker scores (sum of an extended T cell marker gene set) by Pearson correlation, reporting the coefficient of determination (R²) and p-value.

#### Analysis of Patient-Derived Neurospheres Cocultured with T Cells

Single-cell RNA sequencing data from BT333 T cells were processed using the Monocle3^149–151^ framework. Cells were filtered based on UMI counts (≥500 UMIs) and cell hashing quality metrics (>5 hash UMIs per cell, top-to-second-best ratio >2). In total, 98,855 cells were retained, with a mean UMI count of 1,961 per cell. A minimum of 739 cells were retained per dose per drug. Size factors were estimated using estimate_size_factors(), and cell cycle scoring was performed with estimate_cell_cycle() using canonical S-phase and G2M-phase markers. For feature selection, differential gene expression analysis identified T cell induced cancer cell response genes using generalized linear models, with high-value genes selected based on q-value < 0.05 and absolute normalized effect > 0.25. Alternatively, dispersion modeling via estimateDispersionsForCellDataSet() (minimum 100 cells detected) identified the top 5,000 genes by excess dispersion ([dispersion_empirical - dispersion_fit] / dispersion_fit). Dimensionality reduction was performed using preprocess_cds() (num_dim = 20) followed by reduce_dimension() (UMAP with n_neighbors = 20, min_dist = 0.1, max_components = 2, fast_sgd = FALSE). Louvain clustering was applied to PCA space using cluster_cells() (resolution = 7e-5). Aggregate expression scoreswere calculated as log(Σ[normalized_expression] + 1). Statistical comparisons between T cell treatment conditions used two-sample t-tests via ggpubr::stat_compare_means(). Dose-response curves displayed mean ± SE using stat_summary(), with DMSO controls as reference lines.

Multi-resolution Variational Inference (MrVI)^88^ from scvi-tools was applied using the T cell induced cancer cell response genes (q-value < 0.05, |normalized_effect| > 0.25, same list for previous decipher and MrVI analysis). MrVI was configured with sample_key = ‘drug_dose_treatment’ and batch_key = ‘Replicate’, trained for 400 epochs with seed = 0 for reproducibility, and used to compute local sample distances (batch_size = 32). Hierarchical clustering of sample distances was performed using Ward linkage with optimal leaf ordering.

## Lead contact

Further information and requests for resources should be directed to José L. McFaline-Figueroa.

## Materials availability

Materials will be made available upon request.

## Data and code availability

All analyses and code used in this study are included in this published article and its supplementary information. The data discussed in this manuscript will be available at the National Center for Biotechnology Information’s Gene Expression Omnibus (GEO) upon publication. Code relating to data processing and figure generation is available in our GitHub repository (https://github.com/mcfaline-figueroa-lab/sci-Comm-Plex). Any additional information required to reanalyze the data reported in this paper is available from the lead contact upon request.

## Acknowledgements

We thank the Columbia Flow Cytometry Core and Neel H. Shah for providing flow cytometry support. We also would like to thank all members of the McFaline-Figueroa and Azizi labs for helpful suggestions and critical discussion on the study. The authors would like to thank Erin Bush and the JP Sulzberger Columbia Genome Center for next-generation sequencing support.

## Funding

L.S. is supported by a NIH NCI Genome and Epigenome Integrity in Cancer (GEIC) T32 postdoctoral fellowship. E.A. is supported by NIH NCI R00CA230195, NHGRI R01HG012875, and grant number 2022-253560 from the Chan Zuckerberg Initiative DAF, an advised fund of Silicon Valley Community Foundation. J.L.M-F. is supported by NIH NHGRI R35HG011941 and NSF CBET 2146007. The CCTI/HICCC Flow Cytometry Core is supported by NIH grants S10OD020056 and S10OD030282. This research was funded in part through the NIH/NCI Cancer Center Support Grant P30CA013696 and used the Genomics and High Throughput Screening Shared Resource.

## Author Contributions

L.S. and J.L.M-F. conceived the study. L.S. led sample processing and data collection, with assistance from R.M.G., M.V., A.N., M.M., X.S., and H.K.. Additionally, L.S. H.K., E.A., J.L.M-F., adopted TCR CDR3 sequences techniques. L.S., Q.C, J.H., and M.M. performed the data analysis. L.S., J.L.M-F., Q.C., R.R., and M.M. interpreted the data. R.M.G, M.V., and A.S. provided technical support on sci-Plex-GXE protocols, and N.K., J.G, and M.F, provided technical support related to cloning. R.M.G. provided technical support on PDN culturing and drug perturbation. S.W.M. and K.L.L. provided PDN lines and provided advised on neurosphere experiment. J.H. provided analytical support for MrVI analysis, and E.A. advised on data analysis across the study. L.S. H.K., performed the TCR clonotype analysis. Q.C. and R.R. provided support for analysis of GLASS data. L.S., J.L.M.-F., Q.C., A.N., M.M., and H.K. wrote the manuscript.

## Ethics declarations

K.L.L. reports research support to DFCI from Bristol Meyers Squibb (BMS). R.R. is a founder of Genotwin and a member of the Scientific Advisory Board of DiaTech and Flahy. The other authors declare no competing interests.

## Notes

### Summary of Updates

We obtained additional hashing reads for the CRISPRa library and updated the analysis accordingly. Supplementary files updated.

## References

1. Guha, P., Heatherton, K. R., O’Connell, K. P., Alexander, I. S. & Katz, S. C. Assessing the Future of Solid Tumor Immunotherapy. Biomedicines 10, (2022).

2. Batlle, E. & Massagué, J. Transforming Growth Factor-β Signaling in Immunity and Cancer. Immunity 50, (2019).

3. Yuzhalin, A. & Kutikhin, A. Interleukins in Cancer Biology: Their Heterogeneous Role. (Academic Press, 2014).

4. Tufail, M., Jiang, C.-H. & Li, N. Immune evasion in cancer: mechanisms and cutting-edge therapeutic approaches. Signal Transduct Target Ther 10, 227 (2025).

5. Wellenstein, M. D. & de Visser, K. E. Cancer-Cell-Intrinsic Mechanisms Shaping the Tumor Immune Landscape. Immunity 48, 399–416 (2018).

6. Mizuno, S. et al. Immunogenomic pan-cancer landscape reveals immune escape mechanisms and immunoediting histories. Scientific Reports 11, 15713 (2021).

7. Racacho, K. J. et al. The tumor immune microenvironment: implications for cancer immunotherapy, treatment strategies, and monitoring approaches. Frontiers in Immunology 16, 1621812 (2025).

8. Miyaguchi, K. et al. Activated T cell therapy targeting glioblastoma cancer stem cells. Scientific Reports 13, 196 (2023).

9. Wintterle, S. et al. Expression of the B7-Related Molecule B7-H1 by Glioma CellsA Potential Mechanism of Immune Paralysis. Cancer Res 63, 7462–7467 (2003).

10. Freeman, G. J. et al. Engagement of the PD-1 immunoinhibitory receptor by a novel B7 family member leads to negative regulation of lymphocyte activation. J Exp Med 192, 1027–1034 (2000).

11. Lin, H. et al. Understanding the immunosuppressive microenvironment of glioma: mechanistic insights and clinical perspectives. J. Hematol. Oncol. 17, 31 (2024).

12. DiDomenico, J. et al. The immune checkpoint protein PD-L1 induces and maintains regulatory T cells in glioblastoma. Oncoimmunology 7, e1448329 (2018).

13. Saleh, R., Gopal, M., Gilan, O. & Keam, S. Recent Advances and Challenges in Cancer Immunotherapies for Patients with Autoimmune Diseases. (Frontiers Media SA, 2023).

14. Adiputra, P. A. T. et al. Analysis of PD-L1 Expression in Breast Cancer: A Systematic Review and Meta-Analysis in Asian Population. Asian Pac J Cancer Prev 24, 1453–1462 (2023).

15. Zhang, M. et al. The clinicopathological and prognostic significance of PD-L1 expression in gastric cancer: a meta-analysis of 10 studies with 1,901 patients. Sci Rep 6, 37933 (2016).

16. Wang, J. et al. Tumor-associated macrophages and PD-L1 in prostate cancer: a possible key to unlocking immunotherapy efficacy. Aging (Albany NY) 16, 445–465 (2024).

17. Wang, Z. et al. Molecular and clinical characterization of PD-L1 expression at transcriptional level via 976 samples of brain glioma. Oncoimmunology 5, e1196310 (2016).

18. D’Amico, S. et al. Targeting the antigen processing and presentation pathway to overcome resistance to immune checkpoint therapy. Frontiers in Immunology 13, 948297 (2022).

19. Di Tomaso, T. et al. Immunobiological characterization of cancer stem cells isolated from glioblastoma patients. Clin. Cancer Res. 16, 800–813 (2010).

20. Hazini, A., Fisher, K. & Seymour, L. Deregulation of HLA-I in cancer and its central importance for immunotherapy. J Immunother Cancer 9, (2021).

21. Sari, G. & Rock, K. L. Tumor immune evasion through loss of MHC class-I antigen presentation. Curr Opin Immunol 83, 102329 (2023).

22. Hanif, F., Muzaffar, K., Perveen, K., Malhi, S. M. & Simjee, S. U. Glioblastoma Multiforme: A Review of its Epidemiology and Pathogenesis through Clinical Presentation and Treatment. Asian Pac. J. Cancer Prev. 18, 3–9 (2017).

23. Filley, A. C., Henriquez, M. & Dey, M. Recurrent glioma clinical trial, CheckMate-143: the game is not over yet. Oncotarget 8, 91779–91794 (2017).

24. Reardon, D. A., Omuro, A., Brandes, A. A. & Rieger, J. OS10. 3 randomized phase 3 study evaluating the efficacy and safety of nivolumab vs bevacizumab in patients with recurrent glioblastoma: CheckMate 143. Neuro (2017).

25. Reardon, D. A. et al. Effect of Nivolumab vs Bevacizumab in Patients With Recurrent Glioblastoma: The CheckMate 143 Phase 3 Randomized Clinical Trial. JAMA Oncol 6, 1003–1010 (2020).

26. Zhao, J. et al. Immune and genomic correlates of response to anti-PD-1 immunotherapy in glioblastoma. Nat Med 25, 462–469 (2019).

27. Mathewson, N. D. et al. Inhibitory CD161 receptor identified in glioma-infiltrating T cells by single-cell analysis. Cell 184, 1281–1298.e26 (2021).

28. Wang, A. Z. et al. Glioblastoma-infiltrating CD8+ T cells are predominantly a clonally expanded GZMK+ effector population. Cancer Discov. 14, 1106–1131 (2024).

29. Orrego, E. et al. Distribution of tumor-infiltrating immune cells in glioblastoma. CNS Oncol 7, CNS21 (2018).

30. Lin, P. et al. Increased infiltration of CD8 T cells in recurrent glioblastoma patients is a useful biomarker for assessing the response to combined bevacizumab and lomustine therapy. Int Immunopharmacol 97, 107826 (2021).

31. Makarevic, A. et al. Increased Radiation-Associated T-Cell Infiltration in Recurrent IDH-Mutant Glioma. Int J Mol Sci 21, (2020).

32. O’Rourke, D. M. et al. A single dose of peripherally infused EGFRvIII-directed CAR T cells mediates antigen loss and induces adaptive resistance in patients with recurrent glioblastoma. Science Translational Medicine (2017) doi:10.1126/scitranslmed.aaa0984.

33. Liau, L. M. et al. Association of autologous tumor lysate-loaded dendritic cell vaccination with extension of survival among patients with newly diagnosed and recurrent glioblastoma: A phase 3 prospective externally controlled cohort trial: A phase 3 prospective externally controlled cohort trial. JAMA Oncol. 9, 112–121 (2023).

34. Arrieta, V. A. et al. ERK1/2 phosphorylation predicts survival in recurrent glioblastoma following intracerebral and adjuvant PD-1/CTLA-4 immunotherapy: A REMARK-guided analysis. Clin. Cancer Res. 30, 379–388 (2024).

35. Frangieh, C. J. et al. Multimodal pooled Perturb-CITE-seq screens in patient models define mechanisms of cancer immune evasion. Nat. Genet. 53, 332–341 (2021).

36. Larson, R. C. et al. CAR T cell killing requires the IFNγR pathway in solid but not liquid tumours. Nature 604, 563–570 (2022).

37. Ardito, F., Giuliani, M., Perrone, D., Troiano, G. & Lo Muzio, L. The crucial role of protein phosphorylation in cell signaling and its use as targeted therapy (Review). Int. J. Mol. Med. 40, 271–280 (2017).

38. Brea, E. J. et al. Kinase Regulation of Human MHC Class I Molecule Expression on Cancer Cells. Cancer Immunol Res 4, 936–947 (2016).

39. Roskoski, R., Jr. Properties of FDA-approved small molecule protein kinase inhibitors: A 2024 update. Pharmacol. Res. 200, 107059 (2024).

40. Giglio, R. M., et al. A heterogeneous pharmaco-transcriptomic landscape induced by targeting a single oncogenic kinase. bioRxiv (2024) doi:10.1101/2024.04.08.587960.

41. Wang, M. et al. ATR inhibition induces synthetic lethality in mismatch repair-deficient cells and augments immunotherapy. Genes Dev. 37, 929–943 (2023).

42. Cao, X. et al. Small-Molecule Compounds Boost CAR-T Cell Therapy in Hematological Malignancies. Curr. Treat. Options Oncol. 24, 184–211 (2023).

43. McFaline-Figueroa, J. L. et al. Multiplex single-cell chemical genomics reveals the kinase dependence of the response to targeted therapy. Cell Genom 4, 100487 (2024).

44. Li, L., Hou, M. & Fang, S. Application of colony-stimulating factor 3 in determining the prognosis of high-grade gliomas based on magnetic resonance imaging radiomics. Heliyon 9, e15325 (2023).

45. Hu, J. et al. Regulation of tumor immune suppression and cancer cell survival by CXCL1/2 elevation in glioblastoma multiforme. Sci. Adv. 7, eabc2511 (2021).

46. Hayden, M. S. & Ghosh, S. Regulation of NF-κB by TNF family cytokines. Semin. Immunol. 26, 253–266 (2014).

47. Wang, C. et al. Targeting epiregulin in the treatment-damaged tumor microenvironment restrains therapeutic resistance. Oncogene 41, 4941–4959 (2022).

48. Xu, S. et al. CD74 correlated with malignancies and immune microenvironment in gliomas. Front. Mol. Biosci. 8, 706949 (2021).

49. Guo, M. et al. Inhibition of ICAM1 diminishes stemness and enhances antitumor immunity in glioblastoma via β-catenin/PD-L1 signaling. Nat. Commun. 16, 8642 (2025).

50. Purbey, P. K. et al. Opposing tumor-cell-intrinsic and -extrinsic roles of the IRF1 transcription factor in antitumor immunity. Cell Rep. 43, 114289 (2024).

51. Gao, R. et al. Hypoxia-induced MT2A-tetrameric PKM2 interaction maintains PKM2 activity in a copper-ion-dependent manner. Cell Insight 4, 100277 (2025).

52. Krupa, M. M., Pienkowski, T., Tankiewicz-Kwedlo, A. & Lyson, T. Targeting the kynurenine pathway in gliomas: Insights into pathogenesis, therapeutic targets, and clinical advances. Biochim. Biophys. Acta Rev. Cancer 1880, 189343 (2025).

53. Zu, Y. et al. The NAMPT enzyme employs a switch that directly senses AMP/ATP and regulates cellular responses to energy stress. Mol. Cell 85, 2271–2286.e6 (2025).

54. Chang, K.-Y. et al. Specificity protein 1-modulated superoxide dismutase 2 enhances temozolomide resistance in glioblastoma, which is independent of O6-methylguanine-DNA methyltransferase. Redox Biol. 13, 655–664 (2017).

55. Wu, T., Yang, W., Sun, A., Wei, Z. & Lin, Q. The role of CXC chemokines in cancer progression. Cancers (Basel) 15, 167 (2022).

56. Han, F. & Ma, J. Pan-cancer analysis reveals IL32 is a potential prognostic and immunotherapeutic biomarker in cancer. Sci. Rep. 14, 8129 (2024).

57. Tretina, K., Park, E.-S., Maminska, A. & MacMicking, J. D. Interferon-induced guanylate-binding proteins: Guardians of host defense in health and disease. J. Exp. Med. 216, 482–500 (2019).

58. Tinkey, R. A., Smith, B. C., Habean, M. L. & Williams, J. L. BATF2 is a regulator of interferon-γ signaling in astrocytes during neuroinflammation. Cell Rep. 44, 115393 (2025).

59. Camblor, D. G. et al. Genetic variants in the NF-κB signaling pathway (NFKB1, NFKBIA, NFKBIZ) and risk of critical outcome among COVID-19 patients. Hum. Immunol. 83, 613–617 (2022).

60. Yang, Y., Yan, X., Bai, X., Yang, J. & Song, J. Programmed cell death-ligand 2: new insights in cancer. Front. Immunol. 15, 1359532 (2024).

61. Kesarwani, P. et al. Tryptophan metabolism contributes to radiation-induced immune checkpoint reactivation in glioblastoma. Clin. Cancer Res. 24, 3632–3643 (2018).

62. Zhai, L. et al. Infiltrating T Cells Increase IDO1 Expression in Glioblastoma and Contribute to Decreased Patient Survival. Clin Cancer Res 23, 6650–6660 (2017).

63. Wang, D. et al. BIRC3 is a novel driver of therapeutic resistance in Glioblastoma. Sci. Rep. 6, 21710 (2016).

64. Wang, Q. et al. Comprehensive analysis of bulk, single-cell RNA sequencing, and spatial transcriptomics revealed IER3 for predicting malignant progression and immunotherapy efficacy in glioma. Cancer Cell Int. 24, 332 (2024).

65. Yan, Z.-H. et al. Upregulation of DLX2 confers a poor prognosis in glioblastoma patients by inducing a proliferative phenotype. Curr. Mol. Med. 13, 438–445 (2013).

66. Stevanovic, M., Kovacevic-Grujicic, N., Mojsin, M., Milivojevic, M. & Drakulic, D. SOX transcription factors and glioma stem cells: Choosing between stemness and differentiation. World J. Stem Cells 13, 1417–1445 (2021).

67. Zhou, W. et al. An oncohistone-driven H3.3K27M/CREB5/ID1 axis maintains the stemness and malignancy of diffuse intrinsic pontine glioma. Nat. Commun. 16, 3675 (2025).

68. Han, X., Wang, X., Li, H. & Zhang, H. Mechanism of microRNA-431-5p-EPB41L1 interaction in glioblastoma multiforme cells. Arch. Med. Sci. 15, 1555–1564 (2019).

69. Shi, G. & Zhang, Z. Rap2B promotes the proliferation and migration of human glioma cells via activation of the ERK pathway. Oncol. Lett. 21, 314 (2021).

70. Kozole, S. L. & Beningo, K. A. Myosin light chains in the progression of cancer. Cells 13, 2081 (2024).

71. Day, B. W. et al. EphA3 maintains tumorigenicity and is a therapeutic target in glioblastoma multiforme. Cancer Cell 23, 238–248 (2013).

72. Lawson, K. A. et al. Functional genomic landscape of cancer-intrinsic evasion of killing by T cells. Nature 586, 120–126 (2020).

73. Ebrahimi, N. et al. Targeting the NF-κB pathway as a potential regulator of immune checkpoints in cancer immunotherapy. Cellular and Molecular Life Sciences: CMLS 81, 106 (2024).

74. García-Vicente, L. et al. Single-nucleus RNA sequencing reveals a preclinical model for the most common subtype of glioblastoma. Commun Biol 8, 671 (2025).

75. Browaeys, R., Saelens, W. & Saeys, Y. NicheNet: modeling intercellular communication by linking ligands to target genes. Nat Methods 17, 159–162 (2020).

76. Alim, L. F., Keane, C. & Souza-Fonseca-Guimaraes, F. Molecular mechanisms of tumour necrosis factor signalling via TNF receptor 1 and TNF receptor 2 in the tumour microenvironment. Curr. Opin. Immunol. 86, 102409 (2024).

77. Wang, C. Q. et al. An inhibitory role for human CD96 endodomain in T cell anti-tumor responses. Cells 12, 309 (2023).

78. Qian, D. et al. Immunotherapeutic strategies targeting the PVR–TIGIT/CD96/CD226 signaling pathway in glioma treatment. Annals of Medicine 57, 2588717 (2025).

79. Pollack, B. P., Sapkota, B. & Cartee, T. V. Epidermal growth factor receptor inhibition augments the expression of MHC class I and II genes. Clin. Cancer Res. 17, 4400–4413 (2011).

80. Maruyama, T. et al. Inverse correlation of HER2 with MHC class I expression on oesophageal squamous cell carcinoma. British Journal of Cancer 103, 552–559 (2010).

81. Cai, Y.-W. et al. MAP3K1 mutations confer tumor immune heterogeneity in hormone receptor-positive HER2-negative breast cancer. J Clin Invest 135, (2024).

82. Taniguchi, H. et al. ATR inhibition activates cancer cell cGAS/STING-interferon signaling and promotes antitumor immunity in small-cell lung cancer. Sci Adv 10, eado4618 (2024).

83. Zhen, Y. et al. FGFR inhibition blocks NF-ĸB-dependent glucose metabolism and confers metabolic vulnerabilities in cholangiocarcinoma. Nature Communications 15, 3805 (2024).

84. Yu, P. W. et al. Identification of RIP3, a RIP-like kinase that activates apoptosis and NFkappaB. Curr. Biol. 9, 539–542 (1999).

85. Kotliarova, S. et al. Glycogen synthase kinase-3 inhibition induces glioma cell death through c-MYC, nuclear factor-kappaB, and glucose regulation. Cancer Res 68, 6643–6651 (2008).

86. Bhanpattanakul, S. et al. Modulation of MHC expression by interferon-gamma and its influence on PBMC-mediated cytotoxicity in canine mast cell tumour cells. Sci Rep 14, 17837 (2024).

87. Deb, A., Haque, S. J., Mogensen, T., Silverman, R. H. & Williams, B. R. RNA-dependent protein kinase PKR is required for activation of NF-kappa B by IFN-gamma in a STAT1-independent pathway. Journal of immunology (Baltimore, Md. : 1950) 166, (2001).

88. Boyeau, P., et al. Deep generative modeling of sample-level heterogeneity in single-cell genomics. bioRxiv (2022) doi:10.1101/2022.10.04.510898.

89. Ozawa, T. et al. PDGFRA gene rearrangements are frequent genetic events in PDGFRA-amplified glioblastomas. Genes Dev 24, 2205–2218 (2010).

90. Han, P. et al. Genome-Wide CRISPR Screening Identifies JAK1 Deficiency as a Mechanism of T-Cell Resistance. Front Immunol 10, 251 (2019).

91. Atay, C. et al. BRAF Targeting Sensitizes Resistant Melanoma to Cytotoxic T Cells. Clin Cancer Res 25, 2783–2794 (2019).

92. Nazaret, A. et al. Joint representation and visualization of derailed cell states with Decipher. Genome Biol. 26, 219 (2025).

93. Liu, Z. et al. The prognostic and immunological value of guanylate-binding proteins in lower-grade glioma: Potential markers or not? Front. Genet. 12, 651348 (2021).

94. Sadagopan, A., Michelakos, T., Boyiadzis, G., Ferrone, C. & Ferrone, S. Human Leukocyte Antigen Class I Antigen-Processing Machinery Upregulation by Anticancer Therapies in the Era of Checkpoint Inhibitors: A Review. JAMA Oncol 8, 462–473 (2022).

95. Lei, Q. et al. TNIP1-mediated TNF-α/NF-κB signalling cascade sustains glioma cell proliferation. J. Cell. Mol. Med. 24, 530–538 (2020).

96. Wang, Z. et al. The CXCL family contributes to immunosuppressive microenvironment in gliomas and assists in gliomas chemotherapy. Front. Immunol. 12, 731751 (2021).

97. Lee, S. et al. Identification of CXCL8, IL1A, and IL1B as hub genes of therapeutic resistance in glioblastoma multiforme via bioinformatics analysis. Cancer Genomics Proteomics 22, 791–808 (2025).

98. Li, H. et al. Ferritin light chain promotes the reprogramming of glioma immune microenvironment and facilitates glioma progression. Theranostics 13, 3794–3813 (2023).

99. Zhang, Y., Wu, X., Zhu, J., Lu, R. & Ouyang, Y. Knockdown of SLC39A14 inhibits glioma progression by promoting erastin-induced ferroptosis SLC39A14 knockdown inhibits glioma progression. BMC Cancer 23, 1120 (2023).

100. Singh, S. et al. Harnessing ferroptosis to transform glioblastoma therapy and surmount treatment resistance. Cell Death Discov. 11, 448 (2025).

101. Dong, X.-T. et al. Expression and distribution characteristics of human ortholog of mammalian enabled (hMena) in glioma. Chin. J. Cancer Res. 23, 312–316 (2011).

102. Wykosky, J. & Debinski, W. The EphA2 Receptor and EphrinA1 Ligand in Solid Tumors: Function and Therapeutic Targeting. Molecular cancer research : MCR 6, 1795 (2008).

103. Pan, H. et al. High STK40 Expression as an Independent Prognostic Biomarker and Correlated with Immune Infiltrates in Low-Grade Gliomas. International Journal of General Medicine 14, 6389 (2021).

104. Feng, Y. et al. Emerging interleukin-1 receptor-associated kinase 4 (IRAK4) inhibitors or degraders as therapeutic agents for autoimmune diseases and cancer. Acta Pharm. Sin. B. 14, 5091–5105 (2024).

105. Yang, H. et al. STK35 Is Ubiquitinated by NEDD4L and Promotes Glycolysis and Inhibits Apoptosis Through Regulating the AKT Signaling Pathway, Influencing Chemoresistance of Colorectal Cancer. Frontiers in Cell and Developmental Biology 8, 582695 (2020).

106. Lan, Y. et al. STK17B promotes carcinogenesis and metastasis via AKT/GSK-3β/Snail signaling in hepatocellular carcinoma. Cell Death Dis 9, 236 (2018).

107. Vetsika, E.-K. et al. Pediatric gliomas immunity challenges and immunotherapy advances. Cancer Lett. 618, 217640 (2025).

108. Weinstein, J. N. et al. The Cancer Genome Atlas Pan-Cancer analysis project. Nature Genetics 45, 1113–1120 (2013).

109. Kaatsch, P. Effective targeting of PDGFRA-altered high-grade glioma with avapritinib. Cancer Cell 43, 740–756.e8 (2025).

110. Akiyama, T. et al. Stromal Reprogramming through Dual PDGFRα/β Blockade Boosts the Efficacy of Anti-PD-1 Immunotherapy in Fibrotic Tumors. Cancer Res 83, 753–770 (2023).

111. Neftel, C. et al. An Integrative Model of Cellular States, Plasticity, and Genetics for Glioblastoma. Cell 178, (2019).

112. Azizi, E. et al. Single-cell map of diverse immune phenotypes in the breast tumor microenvironment. Cell 174, 1293–1308.e36 (2018).

113. Kirschenbaum, D. et al. Time-resolved single-cell transcriptomics defines immune trajectories in glioblastoma. Cell 187, 149–165.e23 (2024).

114. Li, X., Zhao, L., Li, W., Gao, P. & Zhang, N. HER2-targeting CAR-T cells show highly efficient anti-tumor activity against glioblastoma both in vitro and in vivo. Genes & Immunity 25, 201–208 (2024).

115. Królikowska, A. & Tarnowski, M. CAR-T cells immunotherapy in the treatment of glioblastoma. Cancer Immunology, Immunotherapy : CII 74, 363 (2025).

116. Harly, C., et al. Human γδ T cell sensing of AMPK-dependent metabolic tumor reprogramming through TCR recognition of EphA2. Sci Immunol 6, (2021).

117. Markosyan, N. et al. Tumor cell–intrinsic EPHA2 suppresses antitumor immunity by regulating PTGS2 (COX-2). The Journal of Clinical Investigation 129, 3594 (2019).

118. Binda, E. et al. The EphA2 receptor drives self-renewal and tumorigenicity in stem-like tumor-propagating cells from human glioblastomas. Cancer Cell 22, 765–780 (2012).

119. Shen, W. et al. Prognostic role of EphA2 in various human carcinomas: a meta-analysis of 23 related studies. Growth Factors 32, 247–253 (2014).

120. Qiu, C. et al. Unveiling the therapeutic promise of EphA2 in glioblastoma: a comprehensive review. Discover Oncology 15, 501 (2024).

121. Kim, S. M. et al. Developing CAR-T/NK cells that target EphA2 for non-small cell lung cancer treatment. Front Immunol 16, 1448438 (2025).

122. Chang, F.-L. et al. An auristatin-based antibody-drug conjugate targeting EphA2 in pancreatic cancer treatment. Biochem Biophys Res Commun 688, 149214 (2023).

123. Tandon, M., Vemula, S. V. & Mittal, S. K. Emerging strategies for EphA2 receptor targeting for cancer therapeutics. Expert Opin Ther Targets 15, 31–51 (2011).

124. Salem, A. F., Gambini, L., Udompholkul, P., Baggio, C. & Pellecchia, M. Therapeutic Targeting of Pancreatic Cancer via EphA2 Dimeric Agonistic Agents. Pharmaceuticals (Basel) 13, (2020).

125. Toracchio, L., Carrabotta, M., Mancarella, C., Morrione, A. & Scotlandi, K. EphA2 in Cancer: Molecular Complexity and Therapeutic Opportunities. Int J Mol Sci 25, (2024).

126. Tröster, A. et al. Targeting EPHA2 with Kinase Inhibitors in Colorectal Cancer. ChemMedChem 18, e202300420 (2023).

127. Joseph, R. et al. EphA2- and HDAC-Targeted Combination Therapy in Endometrial Cancer. Int J Mol Sci 25, (2024).

128. Dasari, S. K. et al. Combination of EphA2- and Wee1-Targeted Therapies in Endometrial Cancer. Int J Mol Sci 24, (2023).

129. Wu, Y. et al. MEK inhibition overcomes resistance to EphA2-targeted therapy in uterine cancer. Gynecol Oncol 163, 181–190 (2021).

130. Piccioni, D. et al. Phase II trial of nilotinib in PDGFR-alpha-enriched recurrent high-grade gliomas. Neurooncol Adv 7, vdaf150 (2025).

131. Liu, X. et al. Valtrate, an iridoid compound in Valeriana, elicits anti-glioblastoma activity through inhibition of the PDGFRA/MEK/ERK signaling pathway. J Transl Med 21, 147 (2023).

132. Martinez-Lage, M. et al. Immune landscapes associated with different glioblastoma molecular subtypes. Acta Neuropathol Commun 7, 203 (2019).

133. Heiland, D. H. et al. Comprehensive analysis of PD-L1 expression in glioblastoma multiforme. Oncotarget 8, 42214 (2017).

134. Gai, Q.-J. et al. EPHA2 mediates PDGFA activity and functions together with PDGFRA as prognostic marker and therapeutic target in glioblastoma. Signal Transduct Target Ther 7, 33 (2022).

135. Zhu, G. et al. Microgravity-cultured glioblastoma organoids integrated with microfluidic chip for CAR-γδ T evaluation. *Commun*. Biol. 8, 1791 (2025).

136. Mangena, V. et al. Glioblastoma cortical organoids recapitulate cell-state heterogeneity and intercellular transfer. Cancer Discov. 15, 299–315 (2025).

137. Ronaldson-Bouchard, K. et al. A multi-organ chip with matured tissue niches linked by vascular flow. *Nat*. Biomed. Eng. 6, 351–371 (2022).

138. Shi, L., Lim, J. Y. & Kam, L. C. Substrate stiffness enhances human regulatory T cell induction and metabolism. Biomaterials 292, 121928 (2023).

139. Shi, L., Lim, J. Y. & Kam, L. C. Improving regulatory T cell production through mechanosensing. J. Biomed. Mater. Res. A (2024) doi:10.1002/jbm.a.37702.

140. Srivatsan, S. R. et al. Massively multiplex chemical transcriptomics at single-cell resolution. Science 367, 45–51 (2020).

141. Sziraki, A. et al. A global view of aging and Alzheimer’s pathogenesis-associated cell population dynamics and molecular signatures in human and mouse brains. Nature Genetics 55, 2104–2116 (2023).

142. Cao, J. et al. Comprehensive single-cell transcriptional profiling of a multicellular organism. Science 357, 661–667 (2017).

143. Martin, B. K. et al. Optimized single-nucleus transcriptional profiling by combinatorial indexing. Nat Protoc 18, 188–207 (2023).

144. Hudson, W. H. & Sudmeier, L. J. Localization of T cell clonotypes using the Visium spatial transcriptomics platform. STAR Protoc 3, 101391 (2022).

145. Liu, S. et al. Spatial maps of T cell receptors and transcriptomes reveal distinct immune niches and interactions in the adaptive immune response. Immunity 55, 1940–1952.e5 (2022).

146. Li, S. et al. RNase H-dependent PCR-enabled T-cell receptor sequencing for highly specific and efficient targeted sequencing of T-cell receptor mRNA for single-cell and repertoire analysis. Nat Protoc 14, 2571–2594 (2019).

147. Dobin, A. et al. STAR: ultrafast universal RNA-seq aligner. Bioinformatics 29, 15–21 (2013).

148. Quinlan, A. R. & Hall, I. M. BEDTools: a flexible suite of utilities for comparing genomic features. Bioinformatics 26, 841–842 (2010).

149. Qiu, X. et al. Reversed graph embedding resolves complex single-cell trajectories. Nat Methods 14, 979–982 (2017).

150. Lázár, E. et al. Spatiotemporal gene expression and cellular dynamics of the developing human heart. Nat Genet 57, 2756–2771 (2025).

151. Trapnell, C. et al. The dynamics and regulators of cell fate decisions are revealed by pseudotemporal ordering of single cells. Nat. Biotechnol. 32, 381–386 (2014).

152. Wolock, S. L., Lopez, R. & Klein, A. M. Scrublet: Computational Identification of Cell Doublets in Single-Cell Transcriptomic Data. Cell Syst 8, 281–291.e9 (2019).

153. Traag, V. A., Waltman, L. & van Eck, N. J. From Louvain to Leiden: guaranteeing well-connected communities. Sci Rep 9, 5233 (2019).

154. Wolf, F. A., Angerer, P. & Theis, F. J. SCANPY: large-scale single-cell gene expression data analysis. Genome Biol 19, 15 (2018).

155. Cannoodt, R., et al. SCORPIUS improves trajectory inference and identifies novel modules in dendritic cell development. bioRxiv 079509 (2016) doi:10.1101/079509.

156. GLASS Consortium. Glioma through the looking GLASS: molecular evolution of diffuse gliomas and the Glioma Longitudinal Analysis Consortium. Neuro Oncol 20, 873–884 (2018).

